# Continuous Large Maf Activity Maintains Fast Myofiber Identity and Multicellular Homeostasis in Adult Skeletal Muscle

**DOI:** 10.64898/2026.10.09.758120

**Authors:** Shunya Sadaki, Ryosuke Tsuji, Tatsuya Matsumoto, Takuto Hayashi, Takehito Sugasawa, Takashi Kudo, Naoki Horii, Kei-ichiro Yasunaga, Edgar Jauliac, Pascal Maire, Yusuke Ono, Satoru Takahashi, Ryo Fujita

## Abstract

Fast myofibers are preferentially affected during aging, exhibiting atrophy and shifts toward slower myofiber programs, yet the mechanisms preserving established fast-fiber identity remain unclear. Here, we show that the large Maf transcription factors *Mafa*, *Mafb*, and *Maf* are continuously required to maintain type IIb myofiber identity in adult skeletal muscle. Inducible deletion of large Mafs caused near-complete loss of type IIb fibers, a transition toward type IIx/IIa states, reduced strength, and increased fatigue resistance. RNA-seq and ATAC-seq revealed transcriptional and chromatin accessibility remodeling across the fast myosin locus, characterized by persistent loss of *Myh4* expression and increased accessibility around *Myh1*. Single-nucleus RNA-seq identified transient myonuclei co-expressing multiple myosin isoforms before emergence of a *Myh1*-dominant state, accompanied by remodeling of non-myonuclear cell populations. These changes shared features with aging- and denervation-associated muscle remodeling. Thus, continuous large Maf activity maintains adult fast myofiber identity and coordinates skeletal muscle tissue homeostasis.

## INTRODUCTION

Skeletal muscle accounts for approximately 40% of adult human body mass and is composed of functionally specialized myofibers with distinct contractile and metabolic properties. Adult myofibers are broadly classified as slow-twitch type I or fast-twitch type II myofibers according to their expression of myosin heavy chain (MyHC) isoforms and associated gene programs.^1^ Type I myofibers express *Myh7* (MyHC-I), are enriched in mitochondria, and rely predominantly on oxidative metabolism, supporting sustained and fatigue-resistant contraction. Type II myofibers generate more rapid and forceful contractions and are further classified as type IIa, IIx, or IIb based on the expression of *Myh2*, *Myh1*, or *Myh4*, respectively. These MyHC isoforms differ in their ATPase kinetics and thereby contribute to the characteristic shortening velocities and contractile properties of each myofiber type.^2^ Collectively, these coordinated contractile and metabolic programs establish distinct myofiber identities that enable skeletal muscles to meet diverse functional demands.

Although myofiber identities are established during development, they remain remarkably plastic in adult skeletal muscle, adapting to changes in neural input, mechanical loading, and metabolic demand.^1–3^ Exercise and altered patterns of muscle use can drive coordinated remodeling of contractile and metabolic programs, thereby matching myofiber properties to changing functional requirements.^4^ Myofiber composition is also remodeled during aging and in pathological conditions associated with altered neuromuscular activity.^5–7^ In aging muscle, fast myofibers are particularly susceptible to denervation, atrophy, and loss, and muscle fiber composition progressively shifts toward slower contractile and metabolic phenotypes.^5,8–10^ These changes accompany progressive declines in muscle mass, strength, and metabolic capacity. However, the transcriptional mechanisms that actively maintain an established fast myofiber state in adult muscle remain largely unknown. It is therefore unclear how the failure of such maintenance mechanisms contributes to the erosion of fast myofiber identity during aging and neuromuscular dysfunction.

We previously identified three large Maf transcription factors—MAFA, MAFB, and MAF—as major regulators of fast type IIb myofiber identity in mice.^11^ Large Maf proteins are basic leucine zipper transcription factors that bind Maf recognition elements (MAREs) and control cell type-specific differentiation programs across diverse tissues.^12–15^ Of the four large Maf family members, *Mafa*, *Mafb*, and *Maf* are expressed in skeletal muscle. Simultaneous skeletal muscle-specific deletion of these three genes during development using HSA-Cre resulted in an almost complete loss of type IIb myofibers, accompanied by corresponding increases in type IIx and IIa myofibers.^11^ Conversely, overexpression of individual large Maf factors in adult soleus muscle, which normally contains few type IIb myofibers, induced *Myh4* expression and a broader type IIb myofiber gene program.^11,16^ Large Maf factors similarly activated *MYH4* and broader fast-contractile and glycolytic gene programs in human myotubes,^7,17^ indicating that their function extends beyond the regulation of a single MyHC isoform. Moreover, analyses of human skeletal muscle biopsies revealed positive associations between *MAFA* and *MAF* expression and fast-myofiber abundance.^17^ Together, these findings establish large Maf factors as conserved regulators of fast-myofiber identity rather than merely upstream activators of a single myosin isoform in mice and humans. However, because large Mafs were deleted before type IIb identity was fully established in our previous model,^11^ it remained unclear whether large Maf factors are required only for myofiber specification or continuously maintain fast myofiber identity in adult muscle. Resolving this distinction is essential for understanding how an established myofiber identity is actively maintained in adult muscle and how failure of this maintenance program contributes to fast-myofiber remodeling during aging and disease. The fast *Myh* genes *Myh2*, *Myh1*, and *Myh4* are organized within an approximately 350-kb cluster on mouse chromosome 11, together with the developmental Myh genes *Myh3* and *Myh8* and the extraocular *Myh13* gene.^18^ Within this locus, a 42-kb fast Myh super-enhancer (fMyh-SE), located between *Myh3* and *Myh2*, forms three-dimensional chromatin contacts with transcriptionally active fast Myh promoters.^18,19^ Deletion of the fMyh-SE severely disrupts the expression of the fast Myh genes, demonstrating its central role in coordinating transcription across the locus.^18^ Notably, the fMyh-SE preferentially engages a single fast Myh promoter in individual myonuclei, suggesting that these promoters compete for access to a shared regulatory hub. The fMyh-SE contains several MARE-like sequences^11^, and MAF occupancy has been detected within this region by ChIP-seq.^19^ These observations raise the possibility that large Maf factors stabilize type IIb identity by directing enhancer–promoter interactions toward M*yh4*. However, whether large Maf factors are continuously required to maintain a Myh4-dominant regulatory state in adult muscle, and whether their loss redirects fMyh-SE activity toward *Myh1* and *Myh2* during myofiber-type switching, remain unknown. Here, we generated an adult skeletal muscle-specific inducible triple-knockout (iTKO) model by tamoxifen-induced deletion of *Mafb* and *Maf* in HSA-MerCreMer mice on a *Mafa*-null background. Acute loss of large Maf factors in adult muscle destabilized the *Myh4*-dominant transcriptional state and triggered a progressive transition from type IIb toward type IIx and IIa myofiber identities, as revealed by bulk and single-nucleus transcriptomic analyses and myofiber histology. This transition was accompanied by coordinated remodeling of chromatin accessibility, metabolic programs, and muscle function. Strikingly, disruption of type IIb myofiber identity extended beyond myofibers, eliciting multicellular remodeling of extracellular matrix, angiogenic, and intercellular communication programs that shared features with aging muscle. Together, these findings establish large Maf factors as continuously required regulators of adult type IIb myofiber identity and reveal that disruption of a fast myofiber-intrinsic transcriptional program can reshape the broader skeletal muscle tissue environment.

## RESULTS

### Large Mafs are required to maintain type IIb myofiber identity in adult skeletal muscle

Fast myofibers are preferentially affected by aging and denervation-associated muscle remodeling. To examine whether disruption of the large Maf transcriptional program is associated with the loss of type IIb myofiber identity in adult muscle, we analyzed the expression of fast and developmental myosin heavy-chain (*Myh*) genes and large Maf family members at 1, 3, and 7 days after sciatic nerve transection (SCT). *Myh4* expression remained unchanged during the first 3 days after denervation but declined sharply by day 7 (Figure 1A). At this time point, *Myh2* was markedly increased, whereas *Myh1* remained largely unchanged throughout the observation period (Figure 1B). The developmental myosin genes *Myh3* and *Myh8* progressively increased through day 7 (Figure 1C). Notably, large Maf expression was reduced before the decline in *Myh4*: *Mafa* was already decreased at day 1 and remained low thereafter, whereas *Mafb* and *Maf* decreased from day 3 and remained significantly reduced at day 7 (Figure 1D).

**Figure 1.**
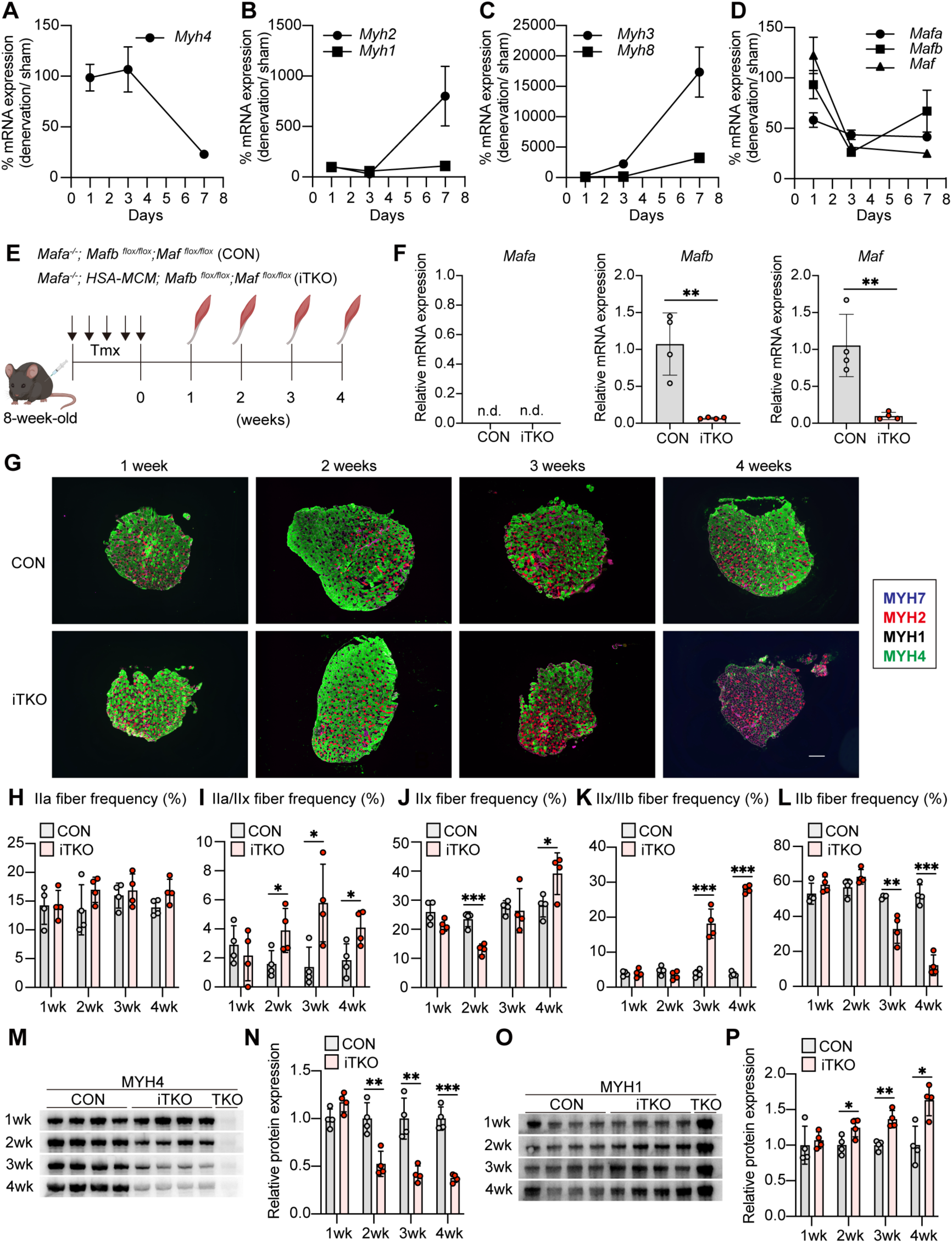
Loss of large Mafs drives adult type IIb-to-IIx myofiber switching. (A–D) Temporal RT-qPCR analysis of *Myh4* (A), *Myh2, Myh1* (B), *Myh3, Myh8* (C), and *large Mafs* (D) expression after sciatic nerve transection (n = 5/group). (E) Experimental scheme for tamoxifen-inducible skeletal muscle-specific large Maf triple-knockout mice. (F) Relative mRNA expression levels of *Mafa*, *Mafb*, and *Maf*, determined by RT-qPCR in gastrocnemius muscles after 1wk-tmx induction (n = 4/group). n.d; not detected. (G) Immunostaining of the myosin heavy chain (MyHC) using myofiber-specific antibodies (MYH7, blue; MYH2, red; MYH4, green) on the EDL muscle. The unstained fibers are considered type IIx myofibers (black). Scale bars, 200 μm. (H-L) Myofiber type composition of IIa (H), IIa/IIx (I), IIx (J), IIx/IIb (K), and IIb (L) in the EDL muscles after knockout induction. (n = 4/group). (M) Western blotting images of MYH4 protein in TA muscles after knockout induction. (N) Quantification of MYH4 protein levels in TA muscles. (n = 4/group). (O) Western blotting images of MYH1 protein in TA muscles after knockout induction. (P) Quantification of MYH1 protein levels in TA muscles. (n = 4/group). P values were calculated using unpaired two-tailed *t*-test; * P < 0.05, ** P < 0.01, *** P < 0.001.

We next examined whether a similar relationship was evident during aging by analyzing a published bulk RNA-seq dataset of gastrocnemius muscles from mice aged 6–27 months.^20^ *Mafa* and *Maf* expression progressively decreased with age, whereas *Mafb* showed no clear age-dependent change (Figures S1A–S1C). *Myh4* expression also declined with age (Figure S1D), while *Myh1* and the developmental myosin genes *Myh3* and *Myh8* increased and *Myh2* remained largely unchanged (Figures S1E–S1H). Thus, both denervation and aging were associated with reduced large Mafs and *Myh4* expression and a shift toward other adult and developmental myosin programs. These shared expression changes raised the possibility that large Maf activity is required not only for the establishment but also for the maintenance of type IIb myofiber identity in adult skeletal muscle.

To directly determine whether large Maf activity is required to maintain type IIb myofiber identity in adult skeletal muscle, we generated inducible skeletal muscle-specific triple-knockout (iTKO) mice lacking *Mafa*, *Mafb*, and *Maf*. iTKO mice carried HSA-MerCreMer (HSA-MCM) and homozygous floxed *Mafb* and *Maf* alleles on a *Mafa*-null background (*Mafa*−/−; *HSA-MerCreMer/+*; *Mafb^flox/flox^; Maf^lox/flox^)* (Figure 1E). Tamoxifen (TMX)-treated *Mafa*-null littermates lacking HSA-MCM served as controls (CON). Eight-week-old mice received tamoxifen for five consecutive days, and skeletal muscles were analyzed at the indicated time points after induction (Figure 1E). At 1 week after TMX administration, the expression of *Mafb* and *Maf* was significantly reduced in skeletal muscle, confirming efficient inducible deletion of the two remaining large Maf genes on the *Mafa*-null background (Figure 1F).

We next examined how myofiber-type composition changed over time in the extensor digitorum longus (EDL) muscle of iTKO mice at 1, 2, 3, and 4 weeks after TMX administration (Figures 1G–1L). Antibodies specific for MyHC I, IIa, and IIb were used to determine myofiber-type composition, and myofibers negative for all three MyHC isoforms were provisionally classified as type IIx myofibers (Figure 1G). At 1 week after induction, myofiber-type composition was comparable between CON and iTKO mice (Figure 1H-1L). By 2 weeks, the proportion of IIa/IIx hybrid myofibers was modestly increased, whereas the abundance of pure type IIb myofibers remained unchanged (Figures 1I and 1L). At 3 weeks, however, the proportion of pure type IIb myofibers was significantly reduced, accompanied by an increase in IIx/IIb hybrid myofibers (Figures 1K and 1L). By 4 weeks, pure type IIb myofibers had markedly declined, accounting for only approximately 20% of total myofibers, whereas pure type IIx and IIx/IIb hybrid myofibers were significantly increased (Figures 1J–1L). To directly confirm that type IIx myofibers increased concomitantly with the loss of type IIb myofibers following large Maf depletion, we performed additional immunostaining using antibodies specific for MyHC IIa, IIx, and IIb (Figure S1I). MyHC IIx immunoreactivity was markedly increased in the EDL muscle of iTKO mice compared with CON mice, confirming that loss of large Maf factors in adult myofibers induces a shift from MYH4-expressing type IIb myofibers toward MYH1-expressing type IIx and hybrid myofiber states.

We further assessed MYH4 and MYH1 protein abundance in the tibialis anterior (TA) muscle by immunoblotting. A reduction in MYH4 and a concomitant increase in MYH1 were already evident at 2 weeks after knockout induction, and both changes became progressively more pronounced through 4 weeks (Figures 1M–1P). Constitutive TKO muscle, included as a control, showed nearly undetectable MYH4 and markedly increased MYH1 abundance, consistent with the previously established loss of the type IIb program in this model.^11^ Together, these results demonstrate that large Maf factors are continuously required to preserve established type IIb myofiber identity in adult skeletal muscle. Their loss initiates a progressive transition from the MYH4-dominant type IIb state toward MYH1-expressing type IIx myofiber states.

### Loss of Large Maf drives oxidative remodeling and myofiber atrophy in fast-dominant muscles

Our previous studies in mouse and human skeletal muscle demonstrated that large Maf factors regulate not only *Myh4* expression but also the glycolytic gene program associated with fast myofiber identity.^11,17^ We therefore investigated whether inducible loss of large Maf factors in adult skeletal muscle shifts the metabolic properties of myofibers from a fast-glycolytic state toward a more oxidative state. To assess oxidative enzyme activity, we performed succinate dehydrogenase (SDH) and NADH-tetrazolium reductase (NADH-TR) staining of EDL muscles at 1, 2, 3, and 4 weeks after TMX administration (Figures 2A–2D). At 1 week after induction, SDH and NADH-TR staining intensities were comparable between CON and iTKO muscles. By 2 weeks, however, both SDH and NADH-TR activities were increased in iTKO muscles, and these differences became progressively more pronounced at 3 and 4 weeks. Thus, inducible loss of large Maf factors in adult skeletal muscle drives progressive oxidative remodeling in fast-dominant muscles, linking the loss of type IIb myofiber identity to a corresponding shift in metabolic properties.

**Figure 2.**
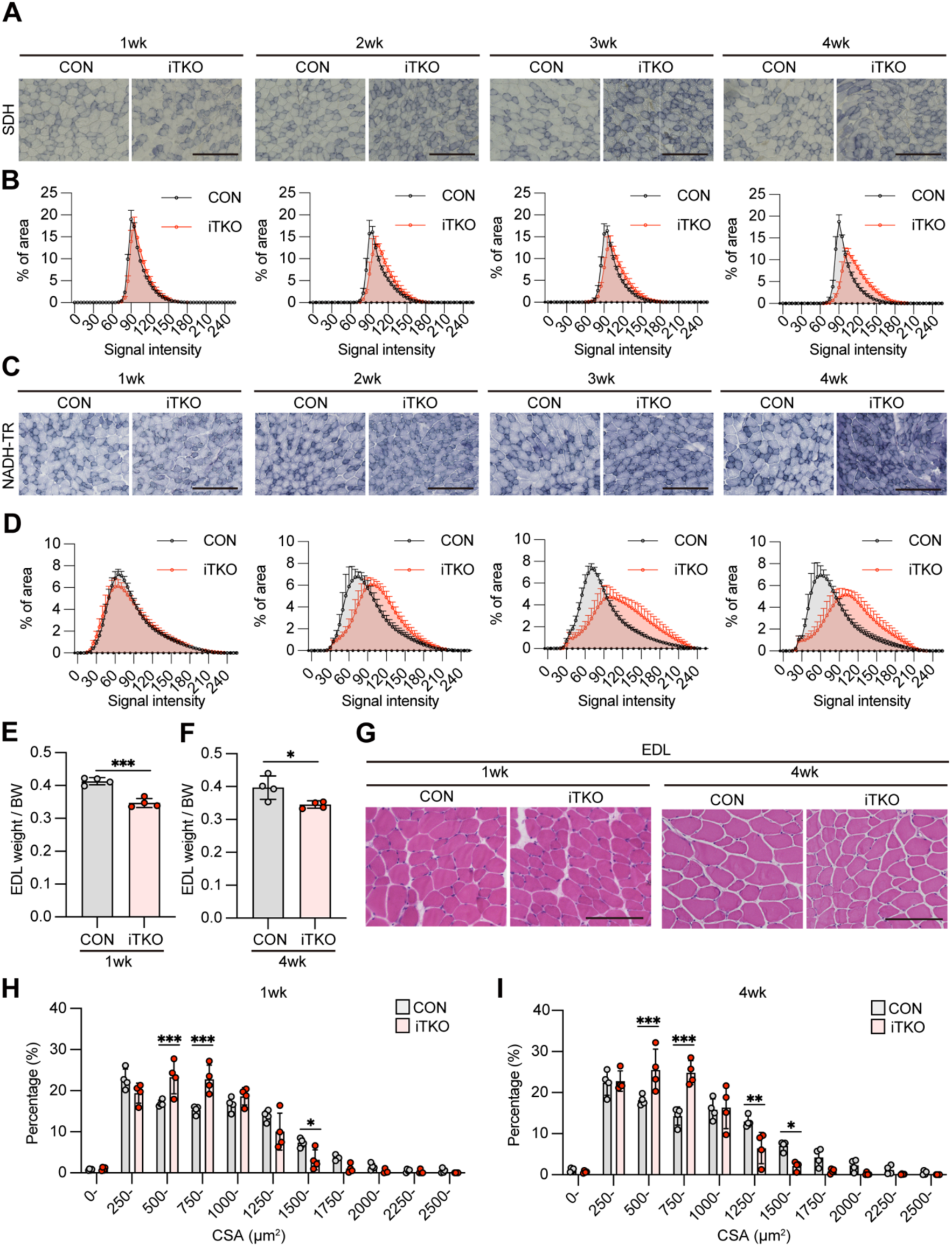
Large Maf loss induces oxidative remodeling and fast-muscle atrophy. (A) SDH staining in the EDL muscle of male CON and iTKO mice. Scale bars, 200 μm. (B) Distribution of SDH signal intensity of the EDL muscle of male CON and iTKO mice (n = 4/group). (C) NADH-TR staining in the EDL muscle of male CON and iTKO mice. Scale bars, 200 μm. (D) Distribution of NADH-TR signal intensity of the EDL muscle of male CON and iTKO mice (n = 4/group). (E, F) EDL muscle weights of male CON and iTKO mice at 1 weeks (E) and 4weeks (F) after tmx induction, normalized to body weight. (n = 4/group). (G) Hematoxylin and eosin staining of EDL muscle cross-sections. Scale bar, 100 μm. (H, I) The CSA distribution in EDL muscles of male CON and iTKO mice at 1 weeks (H) and 4 weeks (I) after tmx induction (n = 4/group).. P values were calculated using unpaired two-tailed *t*-test; *P < 0.05, **P < 0.01, ***P < 0.001.

We previously showed that, among fast myofiber subtypes in mice, type IIb myofibers have the largest cross-sectional area, followed by type IIx and type IIa myofibers. We therefore examined whether loss of large Maf factors affects myofiber size in addition to fast myofiber identity and metabolic properties. Body weight was comparable between CON and iTKO mice before TMX administration but became significantly lower in iTKO mice at 3 and 4 weeks after induction (Figure S2A). Muscle weight normalized to body weight was significantly reduced in fast-dominant hindlimb muscles, including the EDL, gastrocnemius (GAS), and tibialis anterior (TA), in iTKO mice as early as 1 week after induction and remained decreased at 4 weeks (Figures 2E and 2F; Figures S2B and S2C). In contrast, normalized soleus muscle weight was unchanged (Figures S2B and S2C). Hematoxylin and eosin staining revealed no overt histopathological abnormalities, such as centrally nucleated myofibers or inflammatory cell infiltration (Figure 2G). Moreover, TMX-treated HSA-MCM mice showed no differences in normalized muscle weight compared with TMX-treated wild-type (WT) mice at 4 weeks after administration, excluding Cre activity as a primary cause of the reduced muscle weight (Figure S2D).

Consistent with these muscle weight changes, quantification of myofiber cross-sectional area (CSA) revealed a pronounced shift toward smaller myofibers in the EDL of iTKO mice at both 1 and 4 weeks after induction (Figures 2H and 2I). In contrast, CSA was unchanged in the slow myofiber-enriched soleus muscle at either time point (Figures S2E and S2F), indicating that the atrophic phenotype preferentially affected fast myofiber-dominant muscles. Notably, EDL myofiber atrophy was already evident at 1 week, when body weight, myofiber-type composition, and oxidative enzyme activity remained comparable between iTKO and CON mice. Thus, loss of large Maf factors rapidly induces myofiber atrophy before detectable remodeling of myofiber-type composition or metabolic properties, revealing an early requirement for large Maf activity in the maintenance of adult fast myofiber size.

### Loss of large Maf factors disrupts the fast-glycolytic type IIb gene program and activates atrophy-associated transcriptional responses

We next sought to define the transcriptional remodeling induced by loss of large Maf factors in adult skeletal muscle. We performed bulk RNA sequencing of EDL muscles from CON and iTKO mice at 1 and 4 weeks after knockout induction. Principal component analysis (PCA) revealed clear transcriptomic separation between CON and iTKO muscles (Figure 3A). Whereas CON samples collected at 1 and 4 weeks clustered closely together, iTKO samples were clearly separated according to time after induction, indicating that large Maf deficiency elicits dynamic, time-dependent transcriptional remodeling (Figure 3A).

**Figure 3.**
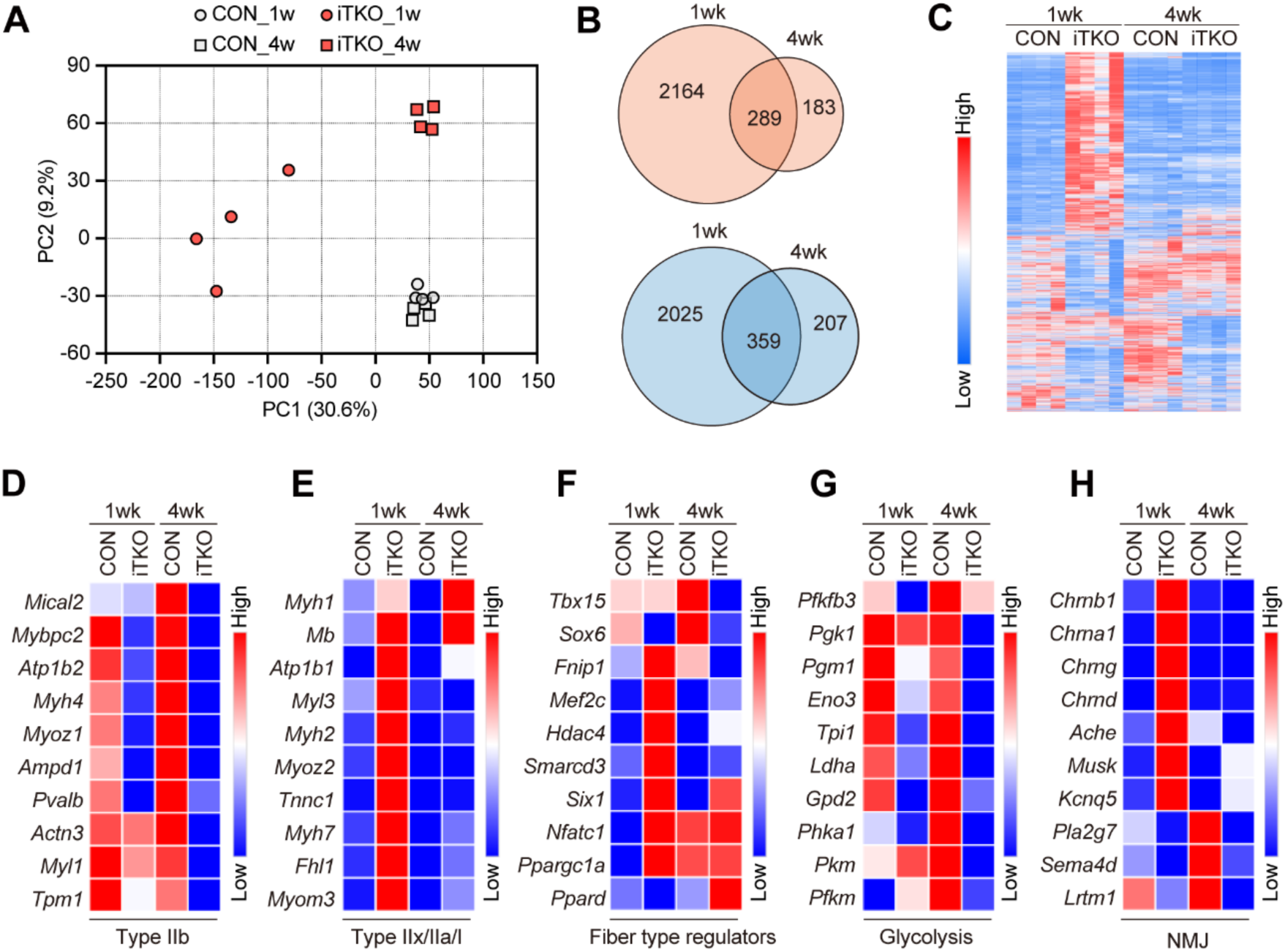
Bulk RNA-seq reveals dynamic transcriptional remodeling after adult large Maf deletion. (A) Principal-component analysis of bulk RNA-seq profiles from EDL muscles of CON and iTKO mice at 1 and 4 weeks after knockout induction. (n = 4 / groups). (B) Venn diagram showing the number of differentially expressed genes (DEGs) upregulated (top) and downregulated (bottom) in EDL muscles of iTKO mice compared with those of CON mice at 1 weeks and 4weeks after knockout induction. (C) Heatmap visualizing differential gene expression in EDL muscles of CON and iTKO mice at 1 weeks and 4 weeks after knockout induction. (n = 4 / groups). (D-H) A heatmap visualizing the expression of type IIb myofiber-associated genes (D), other myofiber type-associated genes (E), glycolytic genes (F), representative myofiber-type regulators (G), and neuromuscular junction-associated genes (H).

Using an FDR cutoff of 0.05, differential expression analysis identified extensive transcriptional changes in iTKO muscles. At 1 week after induction, 4,837 genes were differentially expressed, including 2,453 upregulated and 2,384 downregulated genes. At 4 weeks, 1,138 genes were differentially expressed, comprising 472 upregulated and 566 downregulated genes (Figure 3B). Heatmap analysis further demonstrated distinct transcriptional profiles between CON and iTKO muscles and between the early and later stages after large Maf deletion (Figure 3C). Of these differentially expressed genes, 648 were shared between the 1- and 4-week time points (Figures 3B), indicating that loss of large Maf factors induces both persistent and stage-specific transcriptional responses.

We next performed Gene Ontology (GO) enrichment analysis separately for up- and downregulated DEGs to characterize the transcriptional response to large Maf deletion (Figures S3A–S3F). Among genes altered at both 1 and 4 weeks, upregulated genes were enriched for transcriptional regulation, protein folding, and sarcomere organization, whereas downregulated genes were enriched for muscle contraction, calcium handling, and glycogen metabolism (Figures S3A and S3B). Among 1-week-specific DEGs, upregulated genes were associated with translation and cell division, whereas downregulated genes were enriched for cell adhesion, extracellular matrix, and cytoskeletal organization (Figures S3C and S3D). At 4 weeks, specifically upregulated genes were enriched for mitochondrial energy metabolism and oxidative stress responses, whereas downregulated genes were enriched for endoplasmic reticulum (ER) proteostasis, protein quality control, intracellular protein and ion transport, and innate immune responses (Figures S3E and S3F). These changes suggest that the late response to large Maf deletion extends beyond myofiber metabolic remodeling and may involve broader alterations in muscle tissue homeostasis.

We further examined transcriptional programs associated with myofiber identity. At 1 week after knockout induction, genes associated with the type IIb program were already markedly downregulated (Figure 3D). In contrast, *Myh1*, which encodes the type IIx myosin isoform, showed only a modest trend toward increased expression at 1 week but was robustly upregulated by 4 weeks (Figure 3E). Genes associated with type IIa and type I myofiber programs, including *Myh2* and *Myh7*, as well as established regulators of myofiber-type specification, were transiently induced at 1 week but returned toward CON levels by 4 weeks (Figures 3E and 3F). Glycolytic genes were also broadly downregulated in iTKO muscles, with pronounced suppression evident at 4 weeks after knockout induction (Figure 3G). Importantly, TMX-treated *HSA-MCM* mice showed no significant changes in *Myh* isoform expression compared with TMX-treated WT mice at 4 weeks after administration (Figures S4A–S4E), indicating that the altered *Myh* expression observed in iTKO muscles was not attributable to HSA-MCM activity. Together, these findings demonstrate that loss of large Maf factors disrupts the type IIb transcriptional state and its associated glycolytic gene program, leading to a progressive shift toward a type IIx-like myofiber identity.

We next examined transcriptional programs associated with muscle atrophy (Figures S5A-S5G). Consistent with the rapid reduction in normalized muscle weight and CSA in fast-dominant muscles, heatmap analysis revealed early changes in multiple atrophy-associated genes at 1 week after knockout induction (Figure S5A). Among these genes, *Fbxo32* (Atrogin-1) was significantly induced at 1 week (Figures S5B). In addition, the HDAC4–DACH2–MYOG pathway^21^ was also altered at this time point, with increased *Hdac4* and *Myog* expression and reduced *Dach2* expression (Figure S5C-S5E), together with increased *Runx1* and *Gadd45a* expression (Figure S5F-S5G).^22–24^ These molecular changes have previously been associated with denervation-induced muscle remodeling. Consistent with this, neuromuscular junction-related genes similarly exhibited a transient increase at 1 week (Figure 3H). Together, these findings indicate that the early transcriptional response to loss of large Maf factors partially overlaps with programs activated following denervation.

### Large Maf loss reconfigures the regulatory landscape of the fast myosin heavy-chain gene cluster

The dynamic transcriptional changes following large Maf deletion prompted us to investigate whether loss of large Maf factors also alters the chromatin regulatory landscape underlying fast myofiber identity, including the previously identified fast myosin heavy-chain super-enhancer (fMyh-SE) region.^18^ We performed bulk ATAC sequencing of EDL muscles from CON and iTKO mice at 1 and 4 weeks after knockout induction. TSS enrichment was markedly higher in 1-week iTKO samples than in the other groups (Figure S6A and 6B). We next examined genome-wide changes in chromatin accessibility following large Maf deletion. Differential accessibility analysis revealed widespread chromatin accessibility changes in 1-week iTKO muscles compared with CON muscles (Figure S6C). In contrast, these changes were substantially reduced at 4 weeks (Figure S6D). These findings indicate that large Maf loss induces extensive but largely transient remodeling of the chromatin accessibility landscape in adult skeletal muscle.

To determine how these global chromatin changes relate to direct large Maf target genes, we examined chromatin accessibility and mRNA expression at *Actn3* and *Mybpc2*, two direct large Maf targets preferentially expressed in type IIb myofibers in mice (Figure S6E and S6F).^11^ At both loci, chromatin accessibility was reduced at 1 week and declined further by 4 weeks, approaching the low-accessibility state observed in constitutive developmental TKO muscles generated using *HSA-Cre*. Consistent with these chromatin changes, *Actn3* and *Mybpc2* expression was markedly reduced in iTKO muscles. These observations indicate progressive loss of accessibility at direct large Maf target loci accompanied by silencing of the fast myofiber-associated transcriptional program.

Given the pronounced switch from type IIb to type IIx myofibers following large Maf deletion (Figure 1G-P), we next focused on the chromatin accessibility across the fast *Myh* gene cluster, including the fMyh-SE region (Figure 4A). Chromatin accessibility across the *Myh4*-associated regulatory regions was reduced in iTKO muscles at both 1 and 4 weeks after induction (Figure 4A). In contrast, accessibility around *Myh1* was markedly increased, closely paralleling the transcriptional transition from type IIb to type IIx myofiber identity (Figure 4A). Notably, at 1 week after knockout induction, increased accessibility extended beyond *Myh1* to multiple sites within the fMyh-SE and around *Myh3*, *Myh2*, *Myh8*, and *Myh13*. By 4 weeks, most of these transiently accessible regions were no longer detected, whereas increased accessibility around *Myh1* persisted. These findings reveal a transient and widespread reconfiguration of the fast *Myh* locus during the early phase of myosin heavy-chain remodeling, followed by a more restricted chromatin accessibility pattern centered on *Myh1*.

**Figure 4.**
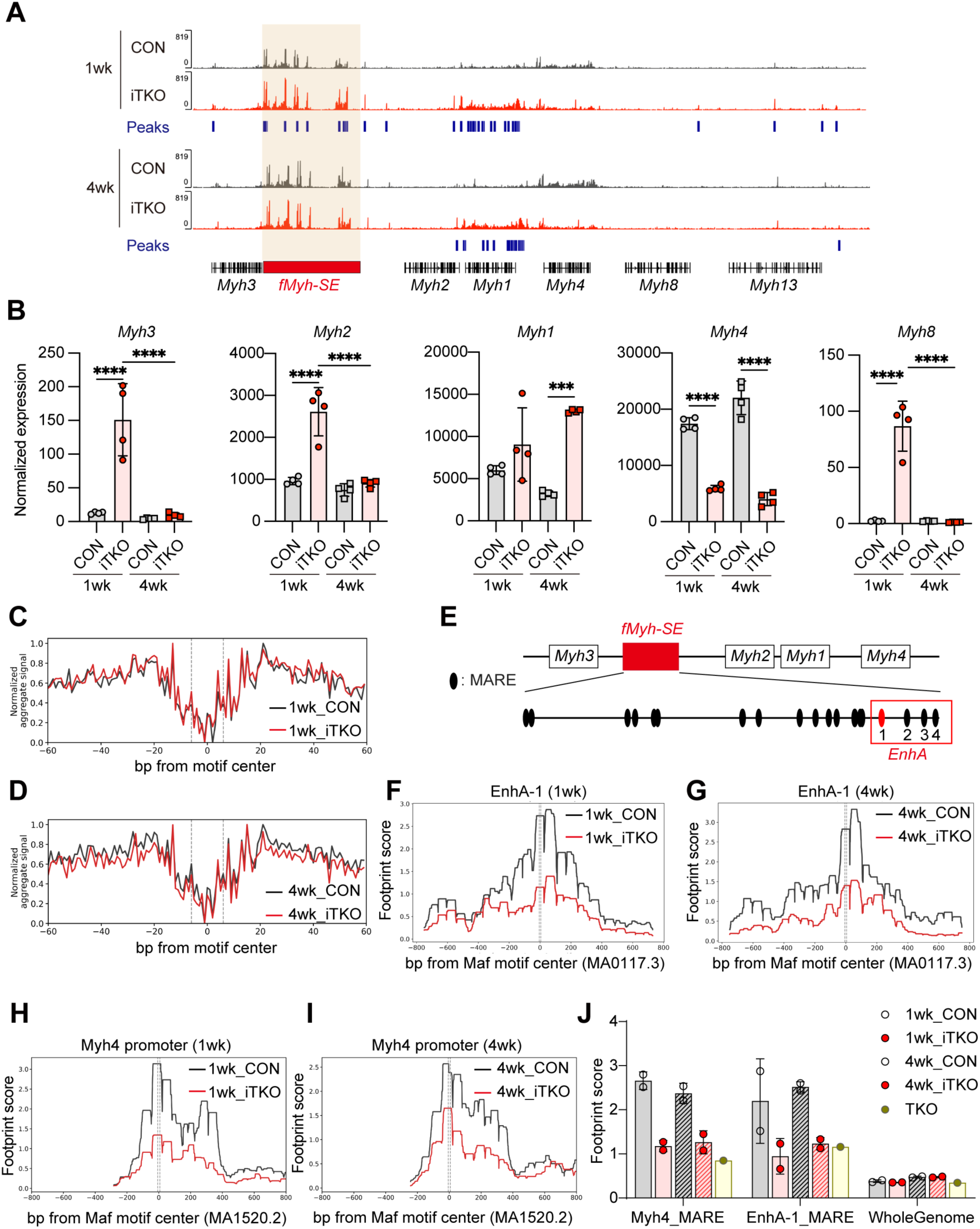
Large Maf deletion remodels chromatin accessibility at type IIb-associated gene loci and the fast *Myh* cluster. (A) Bulk ATAC-seq profiles across the fast *Myh* gene cluster, including the fMyh-SE region, in control and iTKO EDL muscles at 1 and 4 weeks after knockout induction. (B) Bulk RNA-seq expression analysis of fast *Myh* cluster genes in CON and iTKO EDL muscles at 1 and 4 weeks after knockout induction. (C, D) Genome-wide aggregate TOBIAS footprint profiles centered on MAF-family MA1520.2 motifs in control and iTKO EDL muscles at 1 week (C) and 4 weeks (D) after knockout induction. Bias-corrected Tn5 insertion signals are shown relative to the motif center. (E) Schematic of enhancer A within the fMyHC-SE showing the positions of four candidate MAF response elements (MAREs). (F, G) TOBIAS footprint profiles at EnhA-1, the MAF-family motif within enhancer A showing the most prominent reduction in iTKO muscles, at 1 week (F) and 4 weeks (G) after knockout induction. (H, I) TOBIAS footprint profiles at the MAF-family motif within the proximal *Myh4* promoter in control and iTKO EDL muscles at 1 week (H) and 4 weeks (I) after knockout induction. (J) Quantification of TOBIAS footprint scores for genome-wide MAF-family motifs, EnhA-1, and the proximal *Myh4*promoter in control and iTKO muscles at 1 and 4 weeks, with developmental TKO muscles included for comparison. Site-specific footprint scores at EnhA-1 and the *Myh4* promoter were reduced following both adult-inducible and developmental loss of Large Maf factors, whereas changes in genome-wide aggregate MAF footprints were comparatively modest. FDR P values were calculated using edgeR; \**P* < 0.05, \*\*\**P* < 0.001, and \*\*\*\**P* < 0.0001.

Consistent with these changes in chromatin accessibility, *Myh3*, *Myh2*, and *Myh8* expression was transiently increased at 1 week after knockout induction but returned to near-CON levels by 4 weeks (Figure 4B). Interestingly, *Myh1* exhibited a distinct expression pattern, with only a modest trend toward upregulation at 1 week but robust upregulation by 4 weeks (Figure 4B). Conversely, *Myh4*, a direct large Maf target, was sharply downregulated at both time points (Figure 4B). Thus, loss of large Maf factors initially induces broad but transient dysregulation across the fast *Myh* gene cluster, followed by the selective establishment of a *Myh1*-dominant transcriptional state.

To assess MAF-family transcription factor occupancy, we performed TOBIAS footprinting analysis of bulk ATAC-seq data from EDL muscles at 1 and 4 weeks. Genome-wide aggregate profiles showed a clear footprint centered on MAF motifs (MARE, MA1520.2), characterized by reduced Tn5 insertion at the motif center, in both control and iTKO muscles (Figure 4C–4D). Although the aggregate footprint was modestly reduced in iTKO muscles at both time points, the overall genome-wide profiles remained broadly similar between the two groups.

We therefore examined MAF-family footprints at specific regulatory elements within the fast *Myh* locus. Dos Santos et al. previously demonstrated that enhancer A within the fMyHC-SE is functionally important for fast myosin gene regulation.^18^ We identified four candidate MAREs within enhancer A region (EnhA, mm10 chr11:67,138,637–67,145,549) and evaluated TOBIAS footprint profiles individually at each site (Figure 4E). Among these sites, the MAF motif (half MARE, MA0117.3) at EnhA-1 (chr11: 67,139,750–67,139,850, mm10) showed the most prominent and consistent reduction in iTKO muscles at both 1 and 4 weeks (Figures 4F and 4G). Notably, the EnhA-1 site coincided with a reproducible Maf ChIP-seq peak previously identified within enhancer A in adult skeletal muscle,^19^ supporting the interpretation that the reduced ATAC-seq footprint reflects decreased large Mafs occupancy at this regulatory element. MAF-associated footprinting at the proximal *Myh4* promoter was similarly reduced at both time points (Figures 4H and 4I). Quantification of TOBIAS footprint scores confirmed these changes (Figure 4J). At EnhA-1, the mean footprint score decreased from 2.20 in control to 0.95 in iTKO at 1 week and from 2.52 to 1.23 at 4 weeks, corresponding to reductions of 57.0% and 51.1%, respectively. At the proximal *Myh4* promoter, the mean footprint score decreased from 2.66 to 1.18 at 1 week and from 2.52 to 1.23 at 4 weeks, corresponding to reductions of 55.7% and 46.7%, respectively. Footprint scores at both regulatory elements were also reduced in developmental TKO muscles, indicating that diminished MAF-associated footprinting at these sites accompanies both developmental and adult loss of large Maf factors.

Together, these data reveal localized and sustained reductions in MAF-associated footprinting at key regulatory elements within the fast *Myh* locus despite relatively modest changes in genome-wide aggregate MAF footprints. In conjunction with the accompanying chromatin accessibility and transcriptional changes, these findings support a model in which large Maf factors maintain the *Myh4*-dominant regulatory state of adult type IIb myofibers, whereas their loss promotes progressive reconfiguration of the fast *Myh* locus toward a type IIx-associated state.

### Loss of large Mafs promotes transient co-expression of fast *Myh* isoforms in individual myonuclei during myofiber type switching

Bulk RNA-seq and ATAC-seq revealed extensive remodeling of the fast *Myh* gene cluster, including the fMyh-SE, following large Maf depletion. However, these analyses could not distinguish whether the transient upregulation of multiple fast *Myh* genes reflected their co-expression within individual myonuclei or the emergence of transcriptionally distinct myonuclear populations expressing different *Myh* isoforms. To distinguish between these possibilities and define the temporal progression of myosin switching at single-nucleus resolution, we performed single-nucleus RNA sequencing (snRNA-seq) of EDL muscles from CON and iTKO mice at 1 and 4 weeks after knockout induction. Unsupervised clustering of all nuclei identified the major skeletal muscle–associated cell populations across CON and iTKO samples, including myonuclei, muscle stem cells (MuSCs), fibro-adipogenic progenitors (FAPs), smooth muscle cells, Schwann cells, tenocytes, endothelial cells, and B and T cells (Figure 5A). These populations were annotated based on established cell type–specific marker genes (Figure S7A). To examine fast *Myh* regulation during myofiber-type switching, we isolated and re-clustered the myonuclei for further analysis (Figure 5B). Visualization of fast *Myh* transcripts across the resulting myonuclear states enabled us to resolve isoform expression patterns at single-nucleus resolution. Consistent with the bulk RNA-seq and ATAC-seq data, snRNA-seq showed markedly reduced expression of large Maf gene expression in iTKO myonuclei (Figure 5C). Notably, *Myh4* expression was already reduced 1 week after knockout induction, concomitant with increased *Myh1* expression (Figure 5C). Thus, loss of large Maf factors rapidly shifts myonuclei from a type IIb–dominant toward a type IIx–associated transcriptional state.

**Figure 5.**
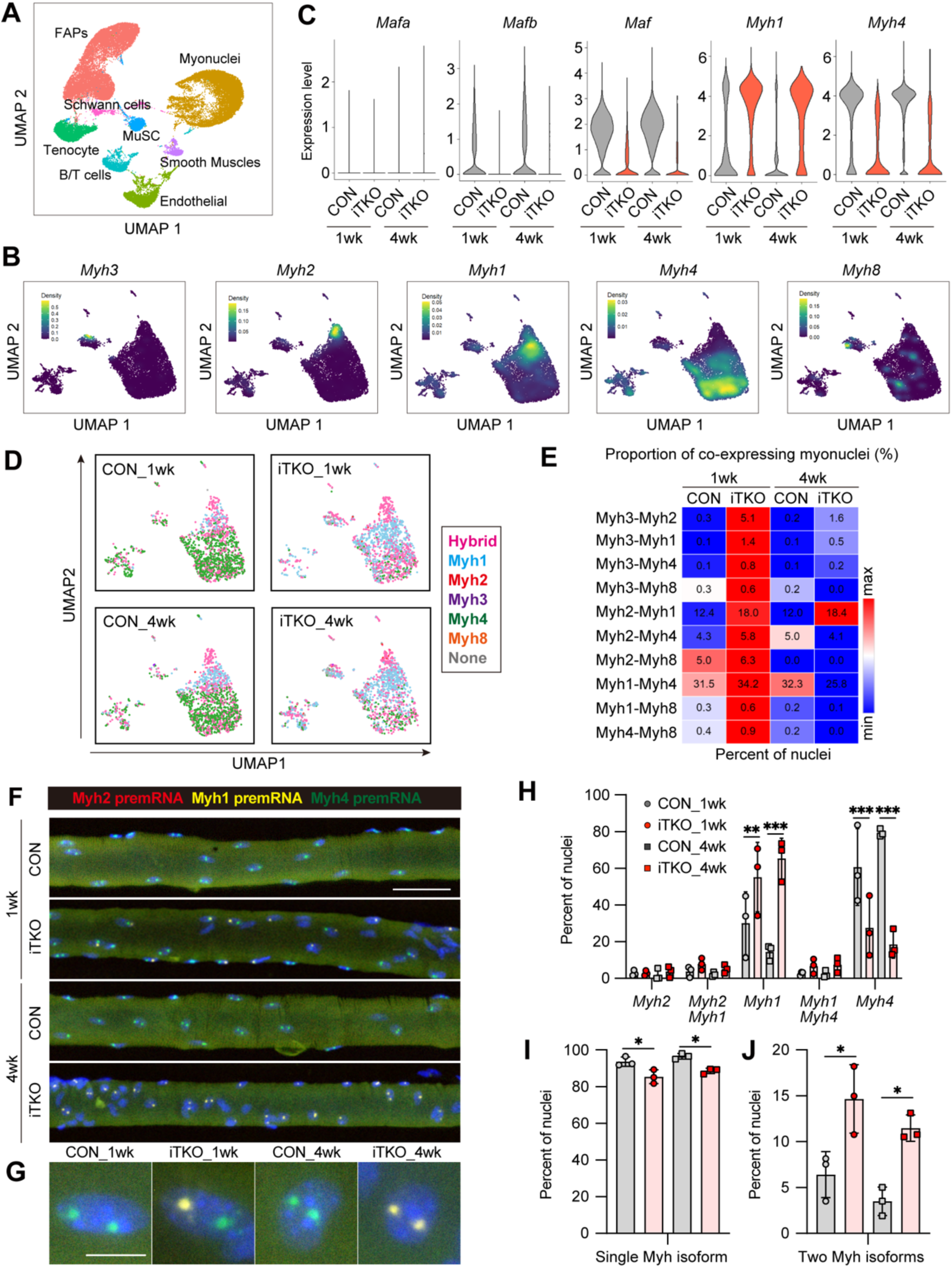
Single-nucleus analysis reveals hybrid fast Myh states after large Maf loss. (A) UMAP visualization and cell-type annotation of snRNA-seq profiles from EDL muscles of control and iTKO mice at 1 and 4 weeks after knockout induction. (B) Re-clustering of myonuclei and distribution of fast Myh gene expression. (C) Violin plots depicting expression patterns of *Mafa, Mafb, Maf, Myh1*, and *Myh4* in myonuclei. (D) UMAP visualization of fast *Myh* expression states in myonuclei from control and iTKO muscles at 1 and 4 weeks after knockout induction. 1,500 myonuclei were randomly sampled from each group for visualization. Hybrid myonuclei co-expressing multiple fast *Myh* genes are increased at 1 week and reduced by 4 weeks, concomitant with the emergence of a *Myh1*-dominant population. (E) Frequencies of pairwise fast *Myh* gene co-expression in individual myonuclei from control and iTKO muscles at 1 and 4 weeks after knockout induction. (F, G) RNAscope analysis of *Myh2*, *Myh1*, and *Myh4* pre-mRNA expression in EDL myonuclei from CON and iTKO mice at 1 and 4 weeks after knockout induction. Representative images are shown in (F), and enlarged views of individual myonuclei are shown in (G). Scale bars; 50 μm in (F) and 10 μm in (G). (H) Quantification of myonuclei expressing each single or pairwise combination of *Myh2*, *Myh1*, and *Myh4* pre-mRNAs. (I,J) Quantification of myonuclei expressing a single fast *Myh* pre-mRNA isoform (I) or two fast *Myh* pre-mRNA isoforms (J). P values were calculated using Tukey’s test (I) and unpaired two-tailed *t*-test (J, K); ^∗^p < 0.05, ^∗∗^p < 0.01, ^∗∗∗^p < 0.001.

To define the broader transcriptional programs accompanying this transition, we performed GO enrichment analysis of DEGs in iTKO myonuclei at 1 and 4 weeks after knockout induction. At both time points, altered genes were enriched for biological processes related to sarcomere organization and muscle contraction (Figures S7B and S7C). At 1 week, additional enrichment was observed for genes involved in the regulation of sarcoplasmic reticulum Ca²⁺ release into the cytosol. By 4 weeks, enriched biological processes included glycogen metabolic processes, glucocorticoid-responsive processes, and regulation of muscle adaptation associated with denervation (Figure S7C). These findings indicate that loss of large Maf factors induces sustained remodeling of contractile, myofiber-type, and metabolic programs in iTKO myonuclei, accompanied by additional time-dependent changes in calcium-handling pathways at 1 week and denervation- and glucocorticoid-related gene sets at 4 weeks.

To determine whether multiple fast *Myh* genes were co-expressed within individual myonuclei, we quantified the number of fast *Myh* genes detected in each myonucleus (Figure S7D). In CON muscles, the majority of myonuclei expressed a single fast *Myh* gene, consistent with previous reports.^25^ Although a minor contribution from residual doublets cannot be fully excluded, iTKO muscles showed a reduction in the proportion of myonuclei expressing a single fast *Myh* gene and a corresponding increase in myonuclei co-expressing multiple fast *Myh* genes (Figure S7D). This increase was primarily attributable to myonuclei co-expressing two fast *Myh* genes and was more pronounced at 1 week than at 4 weeks after knockout induction (Figure S7D). These findings identify a hybrid myonuclear state that is prominent during the early phase of the type IIb-to-IIx transition.

To visualize this transitional state, we mapped fast *Myh* expression patterns onto the myonuclear UMAP (Figure 5D). In CON muscles, myonuclei were dominated by a *Myh4*-expressing population. At 1 week after knockout induction, hybrid myonuclei co-expressing multiple fast *Myh* genes (hybrid) became markedly more abundant, indicating destabilization of the normal *Myh4*-dominant state. By 4 weeks, the abundance of these hybrid myonuclei was reduced, concomitant with expansion of a *Myh1*-dominant population, consistent with progression toward a type IIx-associated myonuclear identity (Figure 5D). We then asked which combinations of fast *Myh* genes were co-expressed within individual myonuclei. Pairwise analysis showed that co-expression frequencies were broadly increased at 1 week after knockout induction across the fast *Myh* gene combinations examined, and these frequencies generally declined by 4 weeks (Figure 5E). Together, these findings indicate that loss of large Maf factors induces a transient hybrid myonuclear state before stabilization of a *Myh1*-dominant type IIx program.

To further characterize these fast *Myh*-co-expressing myonuclei, we identified differentially expressed genes within this population and compared their expression between CON and iTKO muscles (Figure S7E). This analysis identified 527 genes that were altered specifically at 1 week, 331 genes that were commonly altered at both 1 and 4 weeks, and 353 genes that were altered specifically at 4 weeks after knockout induction (Figure S7E). GO enrichment analysis of the 527 genes altered only at 1 week revealed enrichment of biological processes related to transcriptional regulation by chromatin organization and remodeling (Figure S7F).

To investigate whether human skeletal muscle aging is accompanied by analogous changes in *MYH* isoform expression, we re-analyzed published snRNA-seq datasets from young and older individuals.^5^ Aging is associated with denervation, preferential loss of fast myofibers, and remodeling of myofiber-type composition.^26–28^ Moreover, our previous analysis revealed reduced expression of large Maf transcripts in aged human fast myonuclei.^17^ We therefore asked whether aged human myonuclei exhibit *MYH* co-expression patterns reminiscent of those observed in iTKO muscles. In young individuals, approximately 60% of fast myonuclei co-expressed *MYH2* and *MYH1*, and this proportion was further increased in older individuals (Figure S7G). By contrast, myonuclei co-expressing *MYH1* and *MYH8* or *MYH2* and *MYH8* were relatively rare in young muscle but markedly more abundant in older muscle (Figure S7G). Thus, aging is associated with an increased prevalence of myonuclei expressing multiple *MYH* isoforms, including developmental *MYH8*-containing combinations. These findings reveal a partial convergence between aged human myonuclei and large Maf-deficient mouse myonuclei, characterized by increased *MYH* isoform co-expression. Although the relationship between reduced large Maf expression and these age-associated changes remains unresolved, the data suggest that destabilization of fast myonuclear identity may represent a shared feature of large Maf depletion and human skeletal muscle aging.

To independently validate the hybrid myonuclear states identified in iTKO muscles by snRNA-seq, we visualized nascent fast *Myh* transcription within individual myonuclei in mouse EDL myofibers. We performed RNAscope using probes targeting *Myh2*, *Myh1*, and *Myh4* pre-mRNAs (Figures 5F–5J). In single myofibers isolated from CON EDL muscles, the majority of myonuclei with active fast *Myh* transcription expressed *Myh4* pre-mRNA alone (Figures 5F–5H). By contrast, iTKO muscles exhibited a marked increase in the proportion of myonuclei expressing *Myh1* pre-mRNA at both 1 and 4 weeks after knockout induction (Figures 5F–5H). Myonuclei co-expressing *Myh2* and *Myh1* or *Myh1* and *Myh4* pre-mRNAs also tended to be more frequent in iTKO muscles, although neither individual combination reached statistical significance (Figure 5H). The proportion of myonuclei expressing a single fast Myh pre-mRNA was reduced in iTKO muscles (Figure 5I), whereas the combined proportion of myonuclei co-expressing two fast *Myh* pre-mRNAs was significantly increased (Figure 5J). Although the difference between time points did not reach statistical significance, dual-positive myonuclei tended to be more frequent at 1 week than at 4 weeks after knockout induction (Figure 5J). Together, these results provide independent spatial evidence that co-expression of adult fast *Myh* isoforms increases in iTKO myonuclei during myofiber-type switching, supporting the hybrid myonuclear states identified by snRNA-seq.

### Loss of large Maf factors induces multicellular remodeling with aging-associated features

We next asked whether this myofiber-intrinsic remodeling was accompanied by broader changes in the skeletal muscle tissue environment. We first examined the relative representation of each nuclear population across experimental groups (Figure 6A). In iTKO muscles, the relative proportion of myonuclei was reduced at both time points, whereas several non-myonuclear populations were correspondingly overrepresented (Figure 6A). These differences were already detectable 1 week after knockout induction and became more pronounced by 4 weeks. Thus, the myofiber-type transition induced by large Maf depletion was accompanied by progressive shifts in the relative representation of multiple nuclear populations within skeletal muscle.

**Figure 6.**
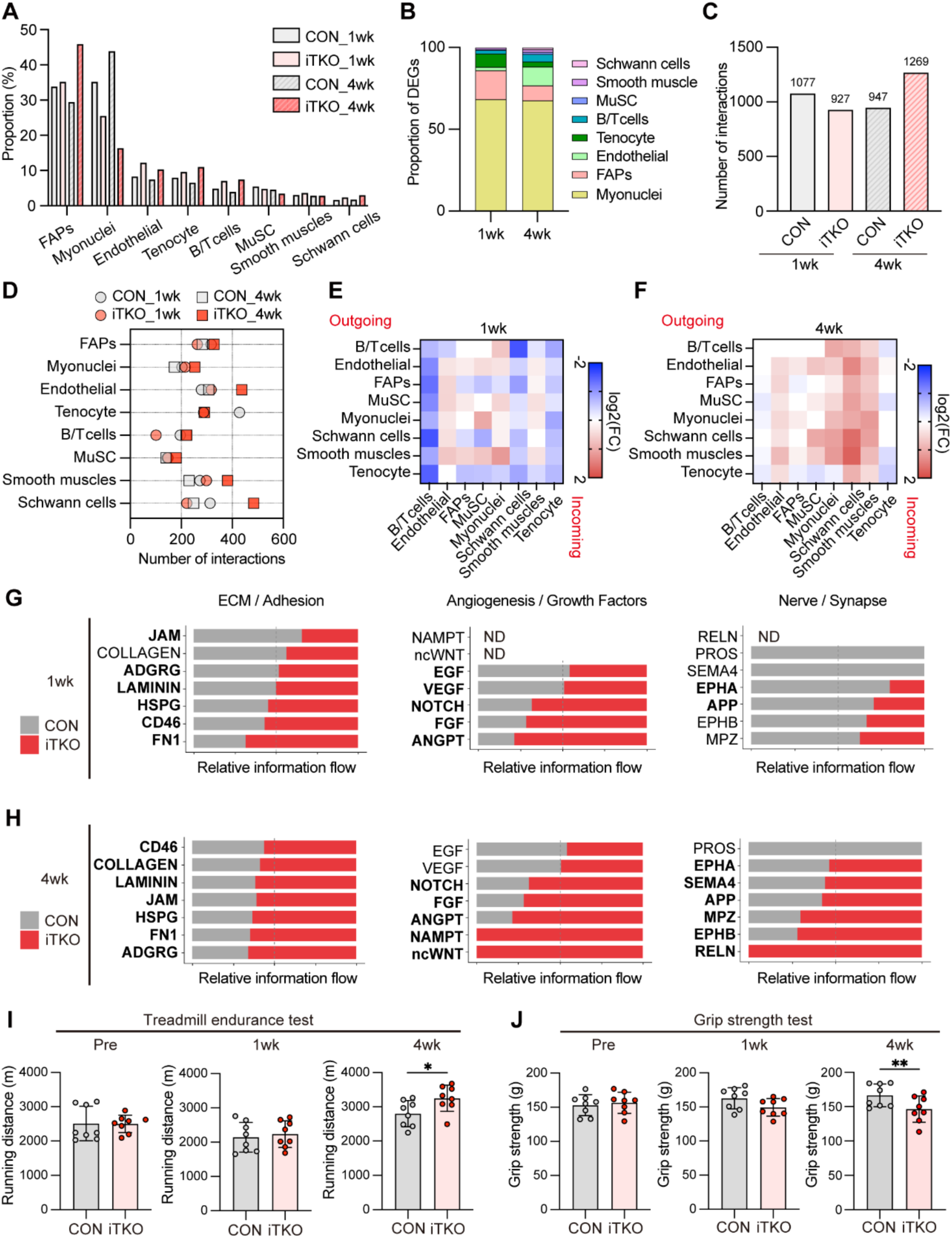
Large Maf depletion induces multicellular and functional remodeling of adult fast muscle. (A) Relative proportions of nuclear populations identified by snRNA-seq in CON and iTKO EDL muscles at 1 and 4 weeks after tmx induction. (B) Distribution of differentially expressed genes across nuclear populations in iTKO muscles at 1 and 4 weeks after knockout induction. (C) Total number of predicted cell–cell interactions inferred by CellChat analysis across experimental groups. (D) Number of predicted CellChat interactions involving each nuclear population in CON and iTKO EDL muscles at 1 and 4 weeks after knockout induction. (E, F) Heatmaps of altered predicted interactions between sender and receiver populations in iTKO muscles compared with CON at 1 week (E) and 4 weeks (F) after knockout induction. (G, H) Predicted CellChat signaling interactions related to ECM/adhesion, angiogenesis/growth factor signaling, and nerve/synapse-associated pathways at 1 week (G) and 4 weeks (H) after knockout induction. (I) Treadmill endurance analysis of CON and iTKO mice before tamoxifen administration and at 1 and 4 weeks after knockout induction (n=8). (J) Grip strength analysis of CON and iTKO mice before tamoxifen administration and at 1 and 4 weeks after knockout induction (n=8). P values were calculated using unpaired two-tailed *t*-test; ^∗^p < 0.05, ^∗∗^p < 0.01.

We next examined the distribution of differentially expressed genes across nuclear populations at 1 and 4 weeks after knockout induction (Figure 6B). At both time points, more than half of all differentially expressed genes (DEGs) were detected in myonuclei, consistent with the primary impact of large Maf depletion on myofiber-intrinsic transcriptional programs (Figure 6B). Nevertheless, a substantial fraction of transcriptional changes occurred in non-myonuclear populations. FAPs accounted for a prominent fraction of the altered genes at 1 week, whereas endothelial cells exhibited more extensive transcriptional remodeling at 4 weeks (Figure 6B). These findings indicate that the consequences of myofiber-specific large Maf depletion extend beyond myonuclei and progressively reshape the transcriptional states of multiple cell populations within the skeletal muscle tissue environment.

We next asked whether the transcriptional remodeling of non-myonuclear populations was accompanied by altered intercellular signaling. We used CellChat to infer ligand–receptor-mediated communication networks across the cell populations identified by snRNA-seq. Quantification of the total number of predicted cell–cell interactions revealed that 4-week iTKO muscles exhibited the largest number of interactions among the groups examined (Figure 6C), suggesting broad expansion of the predicted communication network after myofiber-type remodeling had become established. We next determined which populations contributed to these network-level changes (Figure 6D). At 4 weeks after knockout induction, predicted interactions involving endothelial cells, smooth muscle cells, and Schwann cells were particularly increased. Comparison of the interaction networks between CON and iTKO muscles further revealed distinct time-dependent patterns (Figures 6E and 6F). At 1 week, iTKO muscles showed an overall tendency toward fewer predicted interactions than CON muscles (Figure 6E). By contrast, at 4 weeks, iTKO muscles displayed a broad increase in predicted interactions relative to CON muscles (Figure 6F). Many of these predicted interactions involved myonuclei, suggesting increased signaling potential between remodeled myofibers and surrounding non-myonuclear populations. Schwann cells and smooth muscle cells also exhibited increased predicted connectivity with multiple populations at this time point (Figure 6F).

To obtain a pathway-level overview of these communication changes, we ranked signaling pathways according to differences in relative information flow between CON and iTKO muscles and visualized the top 20 pathways at each time point (Figures S8A and S8B). Based on this analysis, we focused on signaling programs related to three major aspects of skeletal muscle tissue homeostasis: extracellular matrix organization and cell adhesion, angiogenic and growth factor signaling, and nerve- or synapse-associated signaling (Figures 6G and 6H). Although selected predicted signaling interactions within these categories were already increased in 1-week iTKO muscles (Figure 6G), the overall changes were substantially more pronounced at 4 weeks (Figure 6H). Predicted NOTCH, FGF, and FN1 signaling was increased at both time points. In contrast, several signaling pathways, including NAMPT, non-canonical WNT (ncWNT), and RELN, were selectively detected or preferentially increased in 4-week iTKO muscles (Figure 6H). Predicted NAMPT and ncWNT signaling included interactions between myonuclei and endothelial cells, suggesting altered crosstalk between remodeled myofibers and the vascular compartment (Figures S8C and S8D). RELN signaling, by contrast, was associated with predicted communication between myonuclei and Schwann cells (Figure S8E). Previous snRNA-seq studies of aged human skeletal muscle have reported enhanced intercellular signaling involving pathways such as NOTCH, FGF, FN1, and RELN.^5^ The emergence of overlapping signaling programs in iTKO muscles suggests that myofiber-specific large Maf depletion and the resulting loss of type IIb identity in adult muscle partially recapitulate features of aging-associated remodeling within the skeletal muscle tissue environment.

To determine whether the molecular and cellular remodeling induced by large Maf depletion was accompanied by altered muscle function, we assessed treadmill endurance and grip strength before TMX administration (Pre) and at 1 and 4 weeks after knockout induction (Figures 6I and 6J). Treadmill running capacity was comparable between CON and iTKO mice before tamoxifen administration and at 1 week after induction. At 4 weeks, iTKO mice showed a modest but significant increase in running distance (Figure 6I). Grip strength was likewise unchanged before tamoxifen administration and at 1 week after induction but was markedly reduced in iTKO mice at 4 weeks (Figure 6J).

Together, these findings demonstrate that loss of large Maf factors drives broad remodeling of adult fast skeletal muscle, encompassing myofiber-type switching, altered transcriptional states across multiple cell populations, and reorganization of predicted intercellular signaling networks. Functionally, this remodeling was associated with a modest enhancement of endurance capacity and a pronounced loss of muscle strength, consistent with a shift away from a type IIb-dominant, fast-glycolytic muscle phenotype.

## DISCUSSION

Our study identifies large Maf transcription factors as essential regulators of adult fast-glycolytic myofiber identity. Although our previous work established large Maf factors as principal determinants of type IIb myofiber formation during skeletal muscle development,^11^ whether they continue to function in adult myofibers after myofiber identity has been remained unknown. Here, adult myofiber-specific deletion of *Mafb* and *Maf* on a *Mafa*-null background demonstrated that continuous large Maf activity is essential to maintain fast-glycolytic myofiber identity. Loss of large Maf factors induced a progressive transition from MYH4-expressing type IIb myofibers toward MYH1-expressing type IIx through transient hybrid myofiber states. Consistent with this myofiber-type shift, large Maf-deficient muscles exhibited reduced glycolytic features, enhanced oxidative enzyme activity, myofiber atrophy, and multicellular tissue remodeling, accompanied by reduced muscle strength but enhanced endurance capacity. Thus, beyond their developmental role in specifying type IIb myofibers, large Maf factors act as molecular safeguards that preserve fast-glycolytic myofiber identity, fast myofiber size, and tissue homeostasis in adult skeletal muscle.

Fast myofibers are preferentially vulnerable to aging, denervation, and neuromuscular disorders such as amyotrophic lateral sclerosis, and their loss is frequently accompanied by muscle atrophy.^6,9,29^ However, why fast myofibers are selectively susceptible to these perturbations, and which upstream transcriptional changes initiate the destabilization of fast myofiber identity, remain incompletely understood. We found that *Mafa*, *Mafb*, and *Maf* expression declined rapidly after sciatic nerve transection, preceding the marked reduction in *Myh4* expression. This temporal relationship suggests that attenuation of the large Maf program may represent an early event contributing to the loss of the fast-glycolytic myofibers after denervation. Our reanalysis of a mouse gastrocnemius bulk RNA-seq dataset revealed a related expression pattern during aging, characterized by reduced *Mafa*, *Maf*, and *Myh4* expression and increased expression of alternative and developmental myosin genes. Moreover, our previous analysis of human skeletal muscle snRNA-seq datasets showed that *MAF* expression declined with age in fast myonuclei, accompanied by age-associated declines in fast *MYH* gene expression.^5,17^ These observations raise the possibility that attenuation of the large Maf-dependent transcriptional program contributes to the age- and denervation-associated loss of fast myofiber identity. Importantly, our present study demonstrates that adult myofiber-specific depletion of large Maf factors is sufficient to induce rapid myofiber atrophy and extensive remodeling of fast myofiber identity in fast-dominant muscles. Thus, our findings identify loss of large Maf activity as a potential upstream event in denervation- and age-associated fast myofiber remodeling and demonstrate that continuous large Maf activity is required to maintain the fast myofiber program in adult skeletal muscle.

Our integrated transcriptomic, chromatin accessibility, snRNA-seq, and RNAscope analyses suggest that large Maf factors maintain fast myofiber identity by stabilizing a *Myh4*-dominant regulatory state across the fast *Myh* gene cluster. Loss of large Maf factors reduced *Myh4* expression and accessibility at *Myh4*-associated regulatory regions while increasing *Myh1* expression and chromatin accessibility around the *Myh1* locus. It also transiently induced alternative adult and developmental *Myh* programs, including *Myh2*, *Myh3*, and *Myh8*. Thus, the transition from a *Myh4*-dominant to a *Myh1*-dominant state was not a simple reciprocal switch but instead involved a period of reduced myosin isoform selectivity across the *Myh* locus before the establishment of a type IIx-like state. Consistent with this interpretation, single-nucleus analyses revealed that myonuclei co-expressing multiple fast *Myh* genes were most prevalent during the early phase of this transition. These findings indicate that adult myosin switching can proceed through hybrid myonuclear states rather than through the direct replacement of one fixed myonuclear identity by another. Reanalysis of human skeletal muscle snRNA-seq data showed that *MYH*-co-expressing myonuclei were also more frequent in aged human skeletal muscle (Figure S7E). Hybrid myofibers similarly become more prevalent during aging and following denervation, particularly under conditions of incomplete or heterogeneous reinnervation. ^30–33^ These observations suggest that reduced fidelity of *Myh* isoform selection may be a shared feature of large Maf depletion and age- or denervation-associated myofiber remodeling. Large Maf factors may therefore contribute not only to activation of *Myh4* transcription but also to the maintenance of a coherent fast-myosin identity by restricting the emergence of alternative *Myh* programs in adult skeletal muscle.

In addition to destabilizing fast myosin identity, loss of large Maf factors unexpectedly induced rapid myofiber atrophy before overt changes in myofiber-type composition or oxidative enzyme activity became detectable. This atrophic response was prominent in fast-dominant muscles, including the EDL and TA, but was not evident in the slow myofiber-enriched soleus, suggesting that large Maf factors are preferentially required to maintain the mass of adult fast myofibers. The temporal dissociation between atrophy and myofiber-type conversion further indicates that the reduction in myofiber size was not merely a secondary consequence of the transition from type IIb to type IIx identity.

Comparison with our previous developmental TKO model, in which large Mafs were deleted during myofiber formation,^11^ further suggests that the consequences of large Maf loss depend on developmental timing. In the developmental TKO model, fast-dominant muscles contained smaller myofibers without marked induction of atrophy-related genes, suggesting that developing muscles can accommodate the absence of large Maf factors by establishing a smaller, slower myofiber phenotype.^11^ In contrast, acute depletion of large Maf factors from established adult myofibers in iTKO mice triggered rapid atrophy and transient induction of atrogenes, including *Fbxo32*, which encodes Atrogin-1 (Figure S5B). Thus, developmental loss of large Maf factors appears to permit adaptation to an altered myofiber program, whereas their acute loss in adulthood destabilizes an established fast-glycolytic state and elicits a stress-associated atrophic response.

The acute atrophic response following large Maf depletion was accompanied by transient activation of a transcriptional program that partially resembled the response of skeletal muscle to denervation. At 1 week after induction, iTKO muscles showed increased expression of several denervation-responsive genes involved in acetylcholine receptor and neuromuscular signaling,^22^ including *Chrna1*, *Chrnb1*, *Chrnd*, *Chrng*, *Musk*, and *Kcnq5*, whereas their expression largely returned toward CON levels by 4 weeks (Figure 3K). Components of the HDAC4–DACH2–MYOG pathway were also altered during this early phase, and *Runx1* expression was increased in iTKO muscles (Figures S5C–S5F). Given the established roles of class II HDAC–MYOG signaling in denervation-induced *Fbxo32* (Atrogin-1) expression and muscle atrophy,^21,34^ together with the induction of *Runx1* in denervated myonuclei,^22,23,35^ these findings suggest that disruption of the large Maf-dependent fast-muscle program is sufficient to engage a subset of denervation-responsive remodeling pathways.

This model is further supported by our recent collaborative findings showing that denervated type IIb myonuclei acquire increased RUNX1 and AP-1 regulatory activity together with reduced SIX/MAF-associated regulatory signatures.^36^ Large Maf activity may therefore lie at the interface between neural activity and the transcriptional maintenance of fast myofiber integrity. *Maf* expression is reduced when excitation–contraction coupling is disrupted,^19^ and recent work further indicates that fast motor-neuron firing dynamically regulates MAF localization and transcriptional activity.^36^ Conversely, activation of the large Maf-mediated fast myofiber program promotes myofiber hypertrophy^16^ and may attenuate denervation-induced muscle atrophy.^7^ Together, these observations support a model in which neural activity sustains a large Maf-dependent transcriptional program that coordinates *Myh4*-dominant identity with the maintenance of myofiber mass. Thus, large Maf depletion is sufficient to activate a subset of denervation-responsive transcriptional programs in adult fast myofibers without actual denervation.

Beyond myonuclear remodeling, our snRNA-seq revealed extensive changes in the broader muscle tissue environment at 4 weeks after knockout induction, when myofiber-type switching and atrophy were already evident. Although large Maf deletion was restricted to myofibers, transcriptional changes were also detected in non-myonuclear populations, including FAPs, endothelial cells, Schwann cells, and smooth muscle cells. Computational analysis further predicted altered intercellular communication involving ECM-associated FN1 and ADGRG pathways, together with NAMPT and non-canonical WNT signaling between myonuclei and endothelial cells and RELN signaling between myonuclei and Schwann cells. These findings suggest that the consequences of disrupted fast myofiber identity extend beyond myonuclei and are accompanied by multicellular remodeling across stromal, vascular, and neural compartments.

Several of these inferred changes resembled aging-associated features previously described in skeletal muscle, including altered FAP and fibroblast-like cell states, ECM-related transcriptional and signaling programs, and dysregulated growth factor signaling.^5^ Altered NAMPT and ncWNT communication between myonuclei and endothelial cells may reflect remodeling of the metabolic and vascular interface in response to the loss and atrophy of fast myofibers (Figure S8C and S8D). RELN signaling between myonuclei and Schwann cells was among the most prominent computationally inferred pathways, based on relative information flow (Figure S8C and S8D), in both iTKO and aged skeletal muscle.^5^ Although RELN signaling has not been established as a myofiber–Schwann cell communication mechanism in skeletal muscle, its emergence in both iTKO and aged muscles identifies this predicted interaction as a candidate for future investigation. Comparison with the developmental TKO model will be informative in determining whether this multicellular remodeling is specifically triggered by the acute destabilization and atrophy of established adult fast myofibers rather than by the presence of a slower myofiber phenotype itself.

Together with our previous work^11^, the present study clearly establishes large Maf factors as regulators of both the developmental specification and adult maintenance of type IIb myofiber identity in mice. This dual role indicates that type IIb myofiber identity is not a fixed terminal state but an actively maintained transcriptional state that depends on sustained large Maf activity. In human skeletal muscle, *MAFA* and *MAF* expression is associated with fast myofiber gene programs and declines with age in fast myonuclei,^17^ suggesting that attenuation of the large Maf-dependent program may contribute to the preferential vulnerability of fast myofibers in humans. Although it remains to be determined whether reduced large Maf activity directly contributes to fast myofiber loss, atrophy, or functional decline during aging and neuromuscular disease, our findings identify the large Maf-dependent transcriptional network as a central regulator linking fast myofiber identity, myofiber size, muscle function, and multicellular homeostasis in adult. More broadly, our study demonstrates that disruption of a transcriptional program that maintains established myofiber identity can propagate beyond myonuclei, leading to myofiber atrophy, altered muscle function, and multicellular remodeling of adult skeletal muscle.

### Limitations of the study

Several limitations should be considered. First, type IIb myofibers were markedly reduced but not completely eliminated even 4 weeks after tamoxifen-induced large Maf deletion. This residual type IIb identity may reflect incomplete recombination, persistence of pre-existing myosin proteins, or compensation by other fast myofiber regulators such as SIX family transcription factors.^37–39^ Second, the individual contributions of MAFA, MAFB, and MAF remain unresolved. Although our genetic strategy demonstrates the collective requirement for large Maf factors in adult skeletal muscle, individual family members may regulate distinct target genes through interactions with different transcriptional cofactors. Recent work using inducible *Maf* deletion caused myofiber-type shift in specific muscles or regions, including the central region of the TA.^36^ Thus, *Maf* loss alone appears to be largely buffered in many adult fast muscles, likely through functional redundancy among large Maf family members, whereas selected muscles or regions may be more sensitive to *Maf* depletion. Finally, although our ATAC-seq data indicate that loss of large Maf factors alters chromatin accessibility across the fast *Myh* locus and the fMyh-SE, additional large Maf binding and chromatin conformation analyses in adult muscle will be required to define the underlying regulatory architecture and determine whether large Maf factors directly control enhancer–promoter interactions within this locus.

#### RESOURCE AVAILABILITY

### Lead contact

Further information and request for reagents may be directed to and will be fulfilled by the lead contact, Ryo Fujita.

### Materials availability

All unique reagents generated in this study are available from the lead contact in accordance with the relevant material transfer agreements.

## Data and code availability

### Data Availability Statement

- All unique reagents generated in this study are available from the corresponding author, Ryo Fujita, in accordance with the relevant material transfer agreements.
- Bulk RNA-seq data have been deposited in the NCBI Gene Expression Omnibus (GEO) under accession number GSE349174.
- Bulk ATAC-seq data have been deposited in the NCBI Gene Expression Omnibus (GEO) under accession number GSE349176.
- Single-nucleus RNA-seq (snRNA-seq) data have been deposited in the NCBI Gene Expression Omnibus (GEO) under accession number GSE349172.
- This paper does not report original code.
- Any additional information required to reanalyze the data reported in this paper is available from the corresponding author, Ryo Fujita upon request.

## EXPERIMENTAL MODELS AND SUBJECT DETAILS

### Mice

Mice were maintained under specific pathogen-free conditions in the Laboratory Animal Resource Center at the University of Tsukuba, Ibaraki, Japan. *Mafa* knockout mice were generated in a previous study.^15^ Mice harboring the floxed *Mafb* and *Maf* allele were generated using a method described in our previous work.^40,41^ B6.Cg-Tg(ACTA1-cre/Esr1*)2Kesr/J (HSA-MCM, IMSR_JAX:031934) mice were purchased from the Jackson Laboratory (Bar Harbor, ME, USA). *Mafa^-/-^*::*Mafb*^flox/flox^::*Maf*^flox/flox^ mice were crossed with HSA-MCM mice to generate the skeletal muscle specific conditional knockout mice. All experiments were performed in compliance with the relevant Japanese and institutional laws and guidelines and were approved by the University of Tsukuba Animal Ethics Committee (authorization number: 26-218). Tamoxifen (Cayman Chemical) was administered in peanut oil, 10% ethanol by intraperitoneal injections (2.5 mg/day) for 5 days. Denervation was performed in the right hind limb, and a sham operation was performed in the left hind limb. The sciatic nerve was exposed through a skin incision, and a 3–5-mm-long segment of the sciatic nerve was removed. The skin incision was closed using nylon sutures. For the sham operation, identical surgical procedures were performed, except for the nerve transection. Mice were monitored for recovery from anesthesia and then returned to their home cages.

## METHOD DETAILS

### Histological analysis and immunohistochemistry of muscle cryosections

The Achilles tendon side of each hindlimb muscle was vertically mounted on tragacanth gum (Wako, Osaka, Japan) on a cork disc and quickly frozen in isopentane cooled in liquid nitrogen. Frozen tissue sections (8 μm in thickness) were mounted on glass slides.

For hematoxylin/eosin staining, sections were stained with Mayer’s hematoxylin (Wako, Osaka, Japan) for 7 minutes. After washing in warm running tap water (50 ± 5 ℃) for 3 minutes, the sections were stained with eosin (Wako, Osaka, Japan) for 50 seconds, then dehydrated by 2 changes (5 seconds each) of 95% ethanol and 2 changes (5 seconds each) of 100% ethanol (Wako, Osaka, Japan), and finally cleared with 3 changes of xylene (Wako, Osaka, Japan) for 5 minutes each. The sections were mounted with DPX mounting medium (Sigma-Aldrich). All samples were observed with a BIOREVO BZ-X800 microscope (Keyence, Osaka, Japan).

For SDH staining, sections were incubated with a SDH solution (0.2 M sodium succinate, 0.2 M phosphate buffer pH 7.4, 1.2 mM nitro-blue tetrazolium) for 30 minutes at 37 °C. To dehydrate and remove unbound nitro-blue tetrazolium, the sections were then placed into different acetone solutions for 3 seconds in the following order: 30%→60%→90%→60%→30%. After washing with running tap water, the sections were mounted with Glycerol Gelatin (Sigma-Aldrich). All samples were observed within 2 hours of the staining procedure, and intensities were measured using Image J software (National Institute of Health).

For NADH staining, sections were incubated with a NADH-TR solution (1.5 mM NADH and 1.5 mM nitrotetrazolium blue in 200 mM Tris, pH 7.4) for 15 minutes at 37 °C. The sections were then placed into different acetone solutions for 3 seconds in the following order: 30%→60%→90%→60%→30%. After washing with running tap water, the sections were mounted with glycerol gelatin (Sigma-Aldrich). All samples were observed within 2 hours of the staining procedure, and intensities were measured using Image J software (National Institute of Health).

The sections for immunohistochemical analysis were fixed with -20 °C acetone and air-dried. After blocking with 5% goat serum/1% BSA/phosphate-buffered saline (PBS) and M.O.M. blocking reagent (Vector Laboratories, Newark, CA, USA), the samples were incubated with primary antibodies overnight at 4 °C. Antibodies are listed in Key Resource Table. All immunostained samples were visualized using appropriate Alexa Fluor conjugated secondary antibodies (Thermo Fisher Scientific) for 1 h at 37 °C. Tissue sections were mounted with VECTASHIELD Vibrance Antifade Mounting Medium (Vector Laboratories). The proportion of each myofiber type and the CSA of myofibers were assessed using images captured with a BIOREVO BZ-X800 microscope system and hybrid cell count application (Keyence, Osaka, Japan).

### Quantitative analysis of transcripts using reverse transcription PCR

Total RNA was extracted from various tissues and hindlimb muscles using ISOGEN (NIPPON GENE, Chiyoda, Tokyo, Japan). First-strand cDNA was synthesized using ReverTra Ace® qPCR RT Master Mix (Toyobo, Osaka, Japan). Polymerase chain reaction (PCR) was performed using a THUNDERBIRD SYBR qPCR system (Toyobo) and a TP850 Thermal Cycler Dice Real Time System (Takara Bio, Kusatsu, Japan). The primer sequences are listed in Supplementary Table 1. The relative amount of each transcript was normalized to the amount of TATA-box-binding protein (*Tbp)* transcripts.

### Western Blotting

Sample preparation for western blot analysis was performed as previously described, with minor modifications.^42^ The right TA muscle was homogenized in ice-cold cytoplasma extraction buffer (250 mM sucrose, 100 mM potassium chloride, 5 mM EDTA, 20 mM Tris pH 8, protease inhibitor) using a homogenizer and centrifuged at 1,200 × g for 10 min at 4°C. The precipitate was washed in ice-cold buffer containing 0.5% Triton X-100, 175 mM potassium chloride, 2 mM EDTA, 20 mM Tris pH 8, protease inhibitor, centrifuged again at 1,200 × g for 10 min at 4°C, and then dissolved in myofibrillar extraction buffer (50mM DTT, 1% sodium dodecyl sulfate, and 100 mM Tris-HCl pH 6.8, protease inhibitor). After centrifugation at 1,200 × g for 10 min, the supernatant was collected as the myofibrillar protein-rich extract, and heated at 98°C for 10 min.

Protein samples (1 μg) were electrophoresed on 10% sodium dodecyl sulfate-polyacrylamide gels for 30 min at 200 V. Proteins were transferred to polyvinylidene difluoride membranes for 16h at 30V. The membranes were incubated overnight at 4°C with anti-MYH1 (6H1, DSHB) and anti-MYH4 (BF-F3, DSHB), diluted in CanGetSignal Solution A (Toyobo). Then, membranes were washed and incubated at room temperature for 1 h with horseradish peroxidase-conjugated secondary antibodies, diluted in CanGetSignal solution B (Toyobo). Horseradish peroxidase-conjugated secondary antibodies were then visualized by chemiluminescence with iBright 1500 (Thermo Fisher Scientific). The band intensities were normalized to total protein levels.

### Bulk RNA-seq analysis

RNA was extracted from whole frozen EDL tissues using the TRIzoL reagent as indicated in the instructions (Thermo Fisher Scientific). Sequencing libraries of mRNA from three technical replicates were prepared using the NEBNext Ultra II RNA Library Prep Kit for IIumina and the NEBNext Poly(A) mRNA Magnetic Isolation Module (New England Biolabs, pwich, MA, USA), according to the manufacturer’s instructions. The concentration of the library was adjusted to 1 nM and subsequently subjected to denaturation and neutralization. The mRNA-seq libraries were sequenced using NovaSeq X Series 10B Reagent (300 Cycle) Kit (Illumina, San Diego, CA, USA) in the NovaSeq™ X Plus System (Illumina). Sequencing was performed using paired-end reads of 36 bases. The basic information of the NGS run data was assessed using CLC Genomics Workbench 24.0 software (QIAGEN, Hilden, Germany). The read number was approximately 30–48 million per sample, as paired-end reads. Transcripts per million were also obtained using an algorithm “RNA-seq Analysis” as the default setting in CLC Genomics Workbench 24.0 software. Genes with zero counts in any sample were excluded, and differential expression was analyzed using integrated differential expression and pathway analysis. DEGs were extracted among conditions (Control_1wk vs. iTKO_1wk, Control_4wk vs. iTKO_4wk) with a false discovery rate (FDR) of p < 0.05. DAVID Bioinformatics Resources 6.8^43^ was used for gene ontology analysis with P-value < 0.05.

### ATAC-seq analysis

ATAC-seq was performed using EDL muscles from both limbs according to the manufacturer’s instructions (Active Motif, #53150). Nuclei were isolated from muscle tissues, and 100,000 nuclei were used for each assay. The nuclei were lysed in ice-cold ATAC lysis buffer and centrifuged at 500 × g for 5 min at 4°C. The resulting nuclei pellet was resuspended in 50 μL of Tagmentation Master Mix and incubated at 37°C for 30 min with agitation. During this step, accessible chromatin regions were tagged using Tn5 transposase. The tagmented DNA was subsequently purified and PCR-amplified to generate sequencing libraries, which were then subjected to high-throughput sequencing. Sequencing data were analyzed by basepair software (https://www.basepairtech.com/) and were mapped to the mouse genome (mm10), using Bowtie2. ATAC-seq footprinting was performed using TOBIAS^44^. Tn5 sequence bias was corrected using ATACorrect, and footprint scores were calculated using FootprintScores. For large Maf motif analyses, position frequency matrices for MARE (MA1520.2) and half MARE (MA0117.3) were obtained from the JASPAR 2026 CORE vertebrate database. Motif sites within ATAC-accessible regions were identified using TOBIAS BINDetect. Motif-centered aggregate Tn5 profiles were generated from bias-corrected ATAC signals using PlotAggregate, whereas motif-centered footprint-score profiles were generated from TOBIAS FootprintScores bigWig tracks.

### Myonuclear isolation and library preparation

Nuclei were isolated from frozen EDL muscles collected from the CON and iTKO groups at 1 and 4 weeks using the Chromium Nuclei Isolation Kit (10x Genomics). Briefly, frozen muscle tissues were minced, homogenized in Lysis Buffer, and passed through a filtration column. Debris was subsequently removed by centrifugation using the Debris Removal Buffer. The isolated nuclei were washed twice and resuspended in Resuspension Buffer. Nuclei were stained with PI, and their final nuclear numbers were determined under a fluorescence microscope. Nuclei suspensions were adjusted to the appropriate concentration and loaded onto the 10x Chromium system using the Chromium GEM-X Single Cell 3′ Kit v4 (10x Genomics) according to the manufacturer’s instructions. Prepared libraries were sequenced on a NovaSeq X plus platform (Illumina) at Macrogen Japan. Sequenced data was processed using 10x CellRanger (v10.0.0). We used a reference genome built against mouse mm10. The subsequent visualizations, clustering and differential expression tests were performed in R using Seurat (v5.4.0). Quality control on aligned and counted reads was performed keeping cells with >200 and <5000 nFeature RNA and <5% mitochondrial genes. After quality control, 10997, 7154, 13952 and 10576 nuclei from the CON 1-week, iTKO 1-week, CON 4-week, and iTKO 4-week samples, respectively, were retained for downstream analysis. Differentially expressed genes were identified using Seurat FindMarkers, with an adjusted P value < 0.05 and |log2 fold change| > 0.25.

### RNAscope on isolated myofibers

RNAscope on isolated myofibers was performed as previously described using RNAscope® Multiplex Fluorescent Assay V2.^18^ Briefly, EDL muscles were dissected and immediately fixed in 4% PFA at 4 °C during 30 min. Myofibers were dissociated mechanically with small tweezers and fixed onto CREST coated slides (Matsunami Glass) by dehydration at 55 °C during 5 min. Slides were then proceeded according to the manufacturer’s protocol.

### Grip strength test

Forelimb grip strength was determined using a dynamometer (GPM-101B; Melquest, Toyoma, Japan).^11^ The forepaws of mice were allowed to grab a horizontal bar and the tail was slowly pulled back by a researcher. Tension was recorded from the gauge at the time the mouse released its forepaws from the horizontal bar. Measurements were repeated ten times, and mean tension from the ten measurements was used in analyses.

### Treadmill test

Mice were subjected to a low-intensity run-to-exhaustion protocol on a motorized treadmill, as described previously.^45^ Mice were familiarized with the treadmill (Melquest) for 10 min at 10 m/min for two consecutive days. The following day, mice were set to run at 10 m/min for 30 min, 11 m/min for 15 min, and 12 m/min for 15 min. Finally, the speed was incrementally increased by 1 m/min every 10 min until the mouse exhibited exhaustion. The endpoint was reached when the mouse sat on the shock grid at the back of the treadmill and stopped responding to continuous tail brushing for 20 s.

### Quantification and statistical analysis

Graphs are presented as mean ± SEM, as indicated in Figure legends. Significance was calculated using GraphPad Prism software by either Student’s *t*-test or one-way ANOVA with Tukey’s test: *P < 0.05, **P < 0.01, ***P < 0.001, and \*\*\*\**P* < 0.0001, n.d; not detected.

## Acknowledgments

We thank Jiawei Wan of the Laboratory of Clinical Examination and Sports Medicine, Department of Clinical Medicine, Institute of Medicine, University of Tsukuba, for assistance with RNA-seq and ATAC-seq data analysis. We also thank Hiroshi Sakai and Yuki Imai of Ehime University for valuable technical advice and discussions.

## Author Contributions

S.S. and R.F. conceived the study and designed the experiments. S.S., R.T., T.M., T.H., T.S., K.Y., T.K., N.H., Y.O., and R.F. performed the experiments and analyzed the data. E.J., P.M., and S.T. provided scientific guidance and critical input. S.S. and R.F. wrote the manuscript. All authors reviewed and edited the manuscript. R.F. supervised the study.

## Declaration of Generative AI and AI-assisted technologies in the Writing Processes

During the preparation of this work, the authors used ChatGPT for English language editing and the improvement of readability. After using the tool, the authors reviewed and edited the content as needed and take full responsibility for the content of the published article.

## Additional Information

### Supplementary information

Supplementary Information is available for this paper.

