## Supplemental Figures for "Continuous Large Maf Activity Maintains Fast Myofiber Identity and Multicellular Homeostasis in Adult Skeletal Muscle"

Shunya Sadaki *et al.*

**This PDF file includes:**

Figure S1 to S8

Table. S1

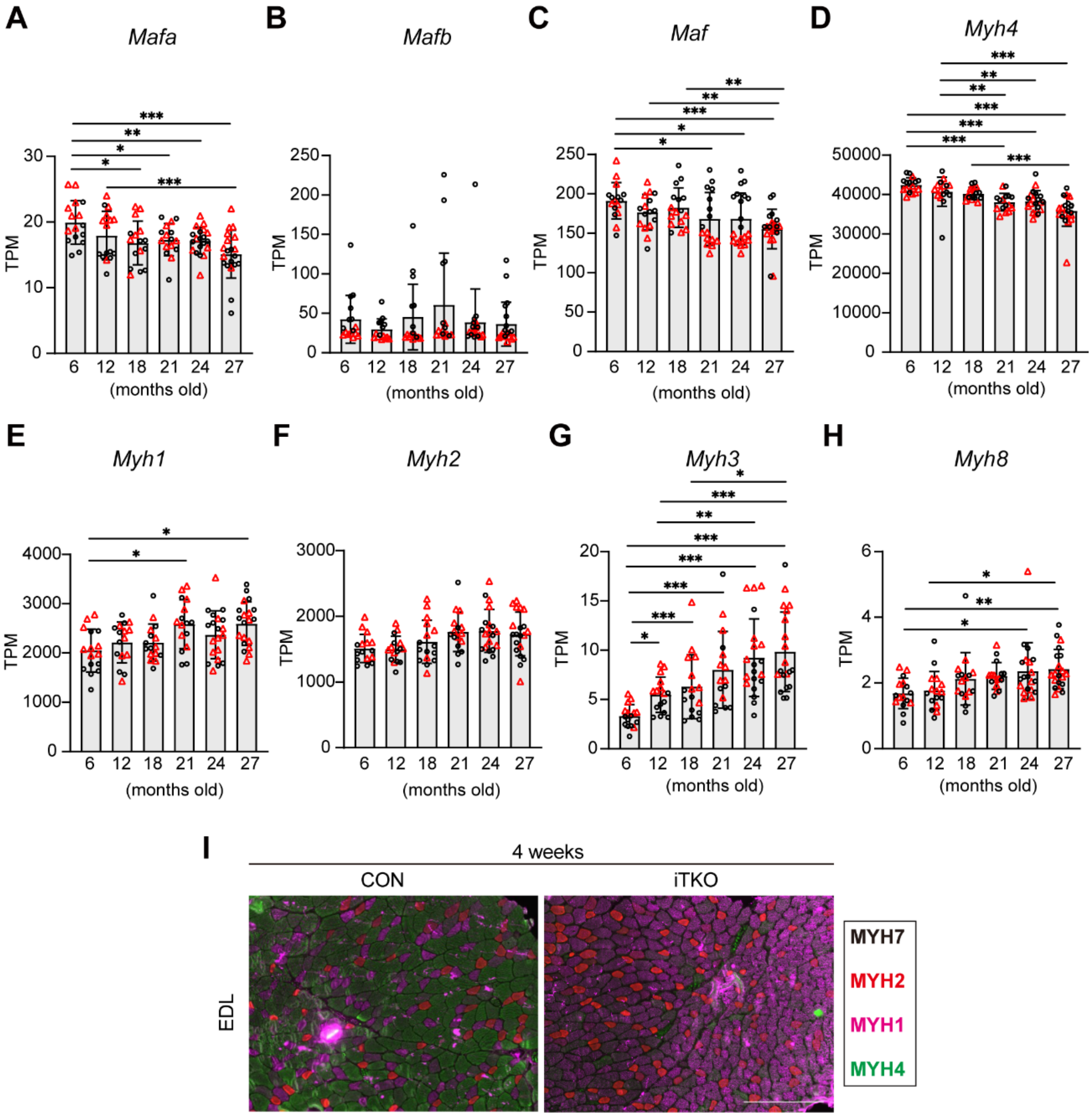


**Figure S1. Loss of large Mafs promotes a transition from type IIb to type IIx myofibers in adult skeletal muscle.**

(A-H) Re-analysis of published RNA-seq data from gastrocnemius muscles of mice aged 6, 12, 18, 21, 24, and 27 months (Shavlakadze et al.). Data are presented as mean ± SEM. Male and female samples are shown in black and red, respectively. FDR P values were calculated using edgeR: $*P < 0.05$, $**P < 0.01$, and $***P < 0.001$. (I) Immunostaining of extensor digitorum longus (EDL) muscle sections from control and iTKO mice at 4 weeks after tamoxifen administration using MyHC IIa-, IIx-, and IIb-specific antibodies. MyHC IIa, red; MyHC IIx, magenta; MyHC IIb, green. Scale bars, 200 μm.

Supplementary Figure 2 (Sadaki et al.)

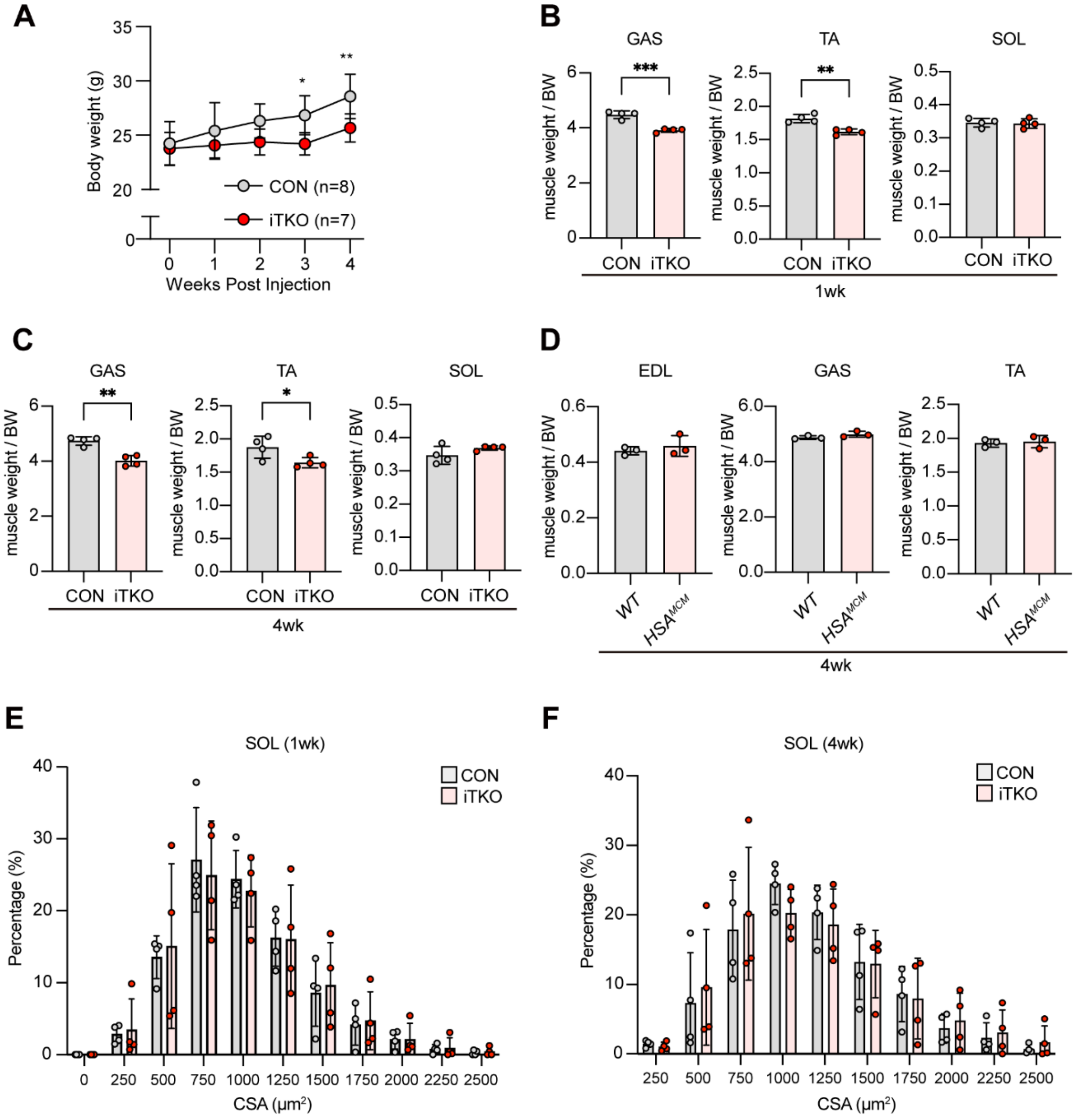


**Figure S2. Loss of large Mafs induces preferential atrophy of fast-dominant muscles without affecting the slow-fiber-enriched soleus.**

(A) Body weight of control and iTKO mice before tamoxifen administration and at 1, 2, 3, and 4 weeks after knockout induction (B and C) Wet weights of the gastrocnemius (GAS), tibialis anterior (TA), and soleus (SOL) muscles normalized to body weight at 1 week and 4 weeks after knockout induction. (D) Wet weights of the indicated muscles normalized to body weight in tamoxifen-treated wild-type (WT) and HSA-MerCreMer (*HSA-MCM*) mice at 4 weeks after tamoxifen administration. (E) Representative cross-sectional area (CSA) distributions and quantification of soleus myofibers in control and iTKO mice at 1 week after knockout induction. (F) Representative CSA distributions and quantification of soleus myofibers in control and iTKO mice at 4 week after knockout induction. Data are presented as mean ± SEM. P values were calculated using unpaired two-tailed *t*-test; * $P < 0.05$, ** $P < 0.01$, *** $P < 0.001$.

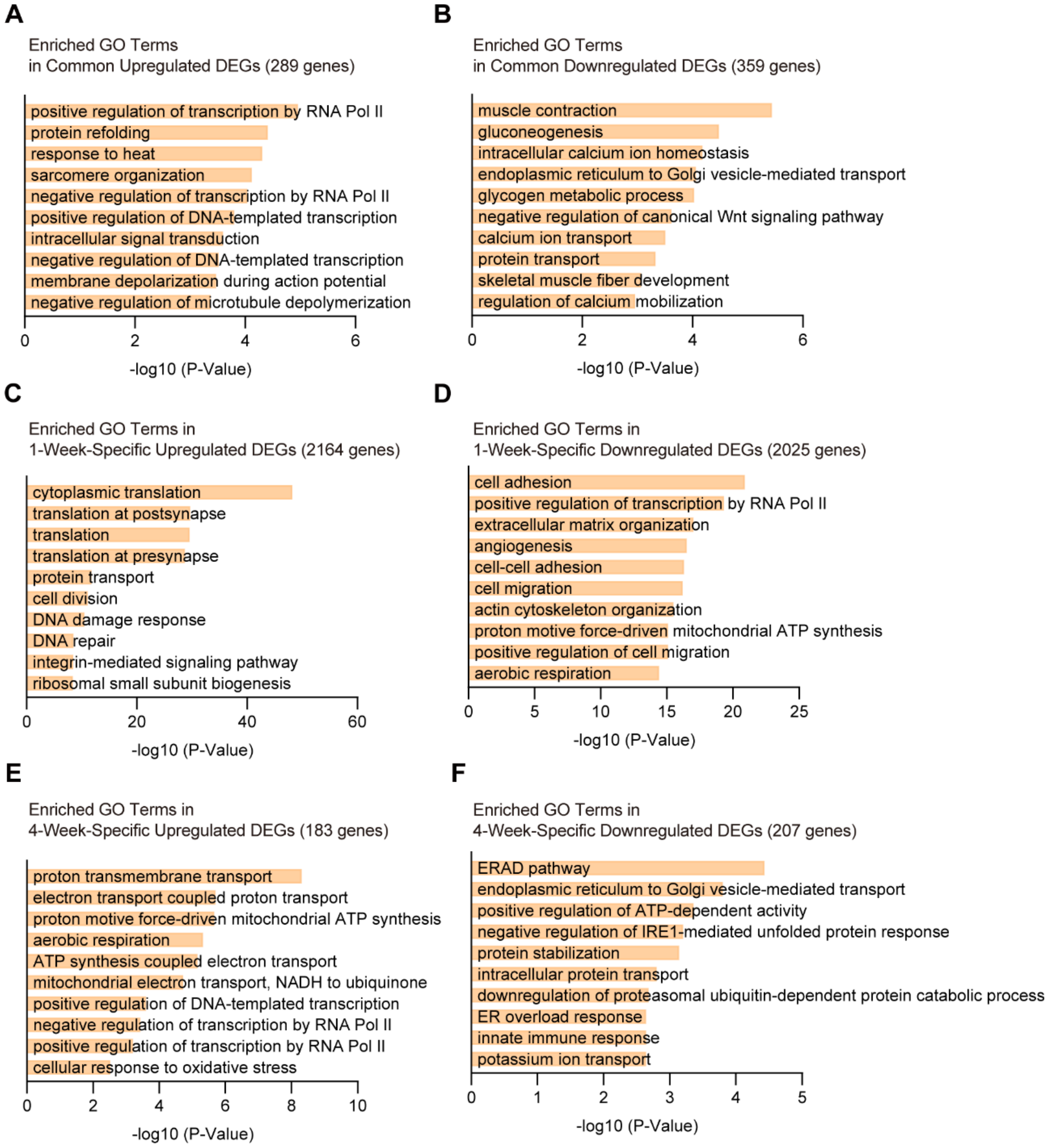


**Figure S3. Gene Ontology analysis of transcriptional changes following adult myofiber-specific large Maf deletion**

(A and B) Gene Ontology (GO) Biological Process terms enriched among genes commonly upregulated (A) or downregulated (B) at 1 and 4 weeks after large Maf deletion. (C and D) GO terms enriched among genes specifically upregulated (C) or downregulated (D) at 1 week. (E and F) GO terms enriched among genes specifically upregulated (E) or downregulated (F) at 4 weeks. Selected significantly enriched GO terms are shown. Ten selected significantly enriched GO terms are shown. Bar length represents the -log10-transformed adjusted *P* value for each GO term.

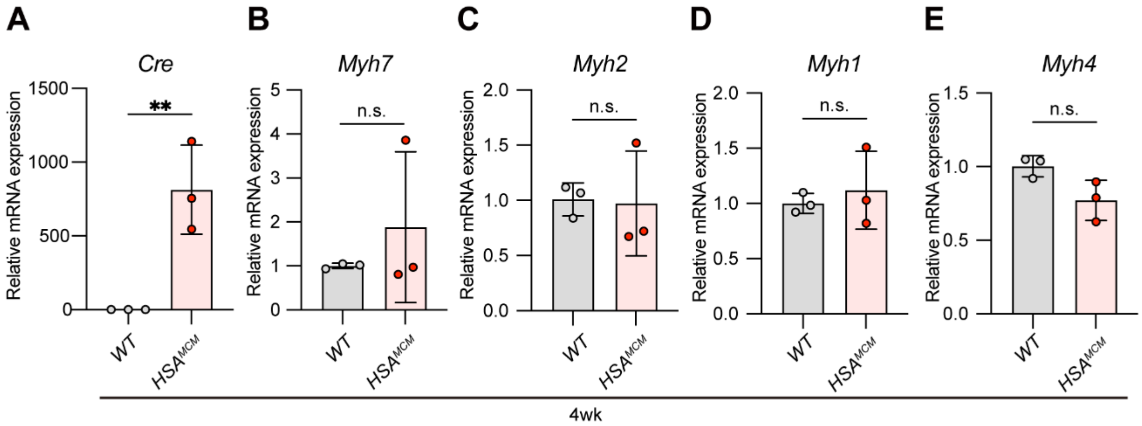


**Figure S4. Tamoxifen-induced HSA-MerCreMer activity does not alter Myh isoform expression in skeletal muscle.**

(A–E) RT-qPCR analysis of Cre (A), *Myh7* (B), *Myh2* (C), *Myh1* (D), and *Myh4* (E) expression in skeletal muscle from tamoxifen-treated WT and *HSA-MCM* mice at 4 weeks after tamoxifen administration. Gene expression levels were normalized to Tbp and are presented relative to those in WT mice. Data are presented as mean ± SEM. P values were calculated using an unpaired two-tailed t-test, **P < 0.01 ; ns, not significant.

Supplementary Figure 5 (Sadaki et al.)

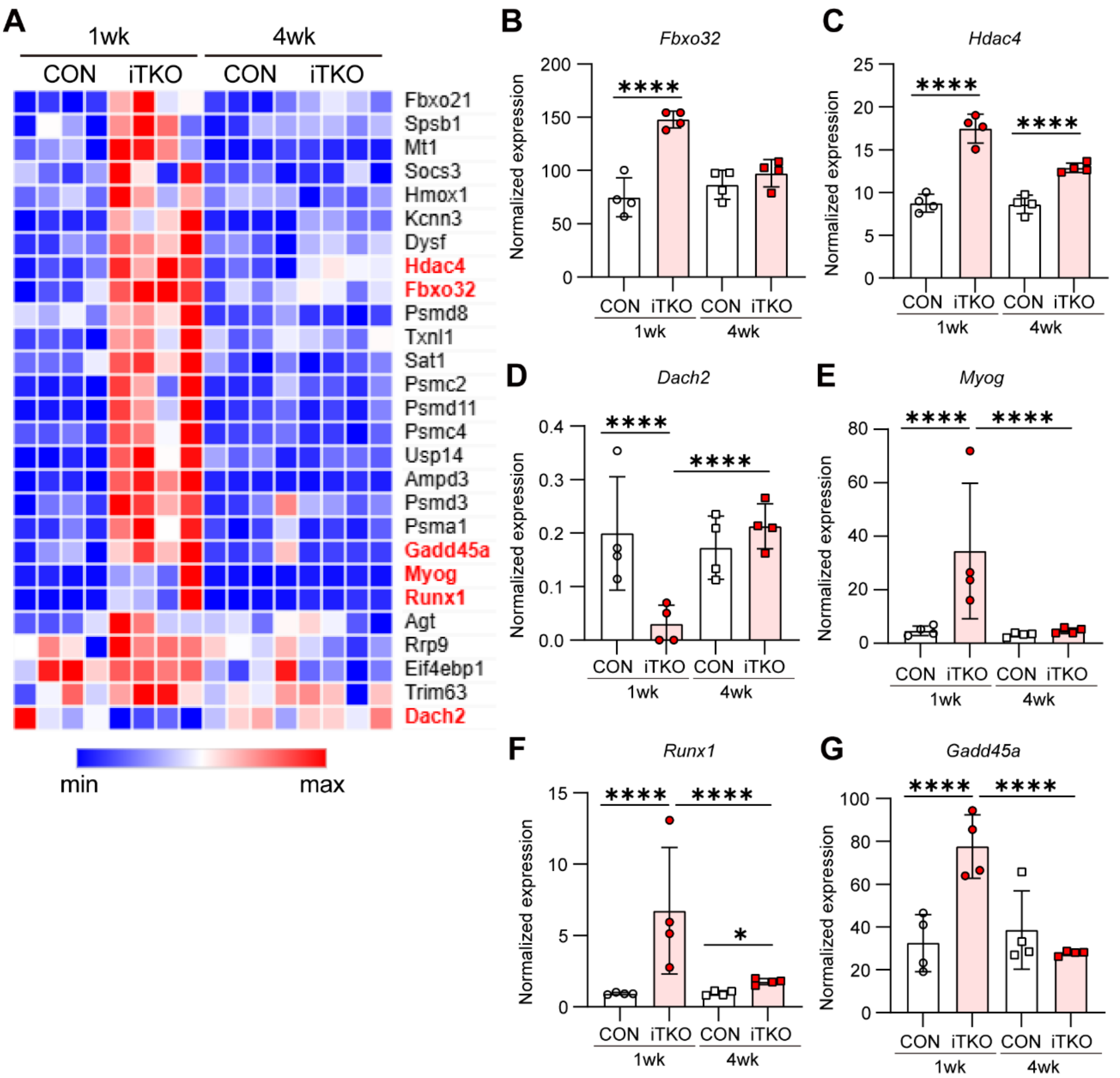


**Figure S5. Loss of large Mafs induces an atrophy-associated transcriptional program in EDL muscles that partially overlaps with denervation-induced remodeling.**

(A) Heatmap showing the expression of atrophy-associated genes significantly altered at 1 week after knockout induction. (B–G) Normalized RNA-seq expression of *Fbxo32* (B), *Hdac4* (C), *Dach2* (D), *Myog* (E), *Runx1* (F), and *Gadd45a* (G) in EDL muscles from control and iTKO mice at 1 and 4 weeks after knockout induction. FDR P values were calculated using edgeR; *$P < 0.05$, and ****$P < 0.0001$.

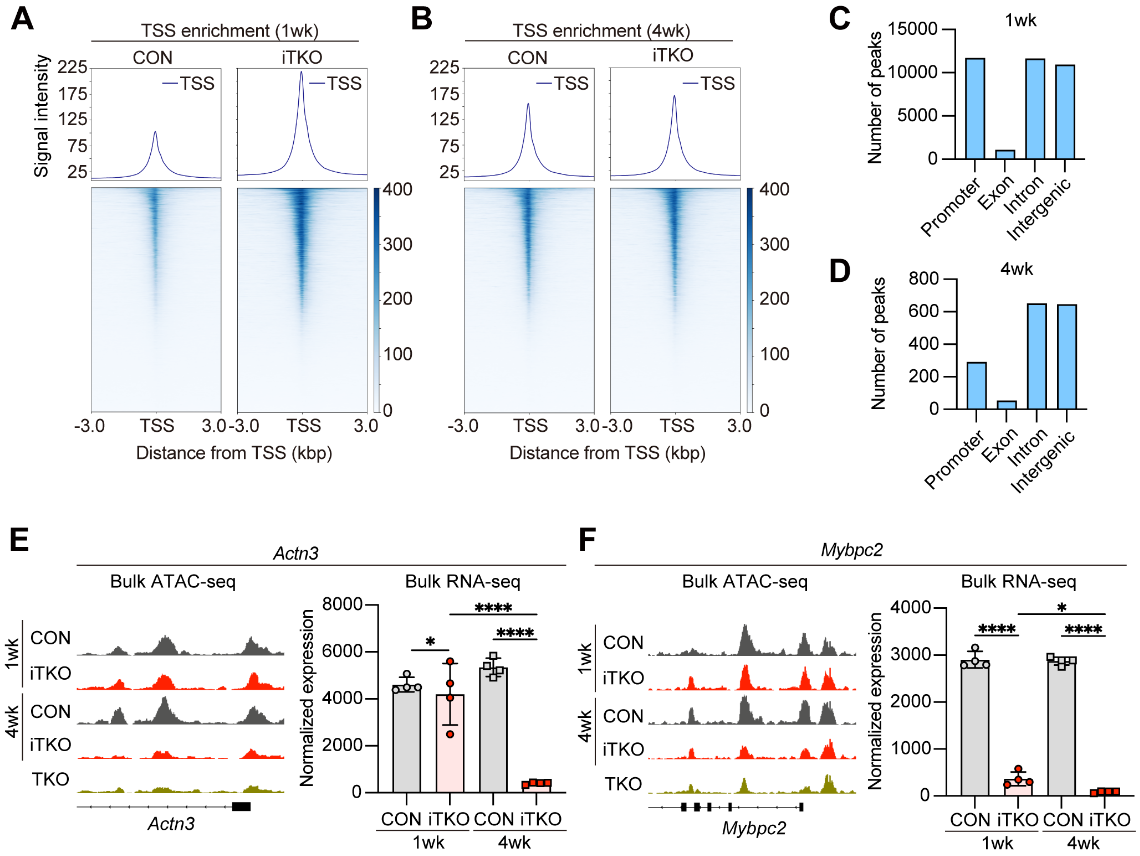


**Figure S6. Large Maf deletion alters chromatin accessibility at fast myofiber-associated genes**
(A, B) ATAC-seq transcription start site (TSS) enrichment profiles and heatmaps in EDL muscles from control and iTKO mice at 1 week (A) and 4 weeks (B) after knockout induction. (C, D) Genomic annotation of differentially accessible ATAC-seq peaks in iTKO EDL muscles at 1 week (C) and 4 weeks (D) after knockout induction. (E, F) Bulk ATAC-seq profiles and bulk RNA-seq expression analysis of the large Maf target genes *Actn3* (E) and *Mybpc2* (F).

Supplementary Figure 7 (Sadaki et al.)

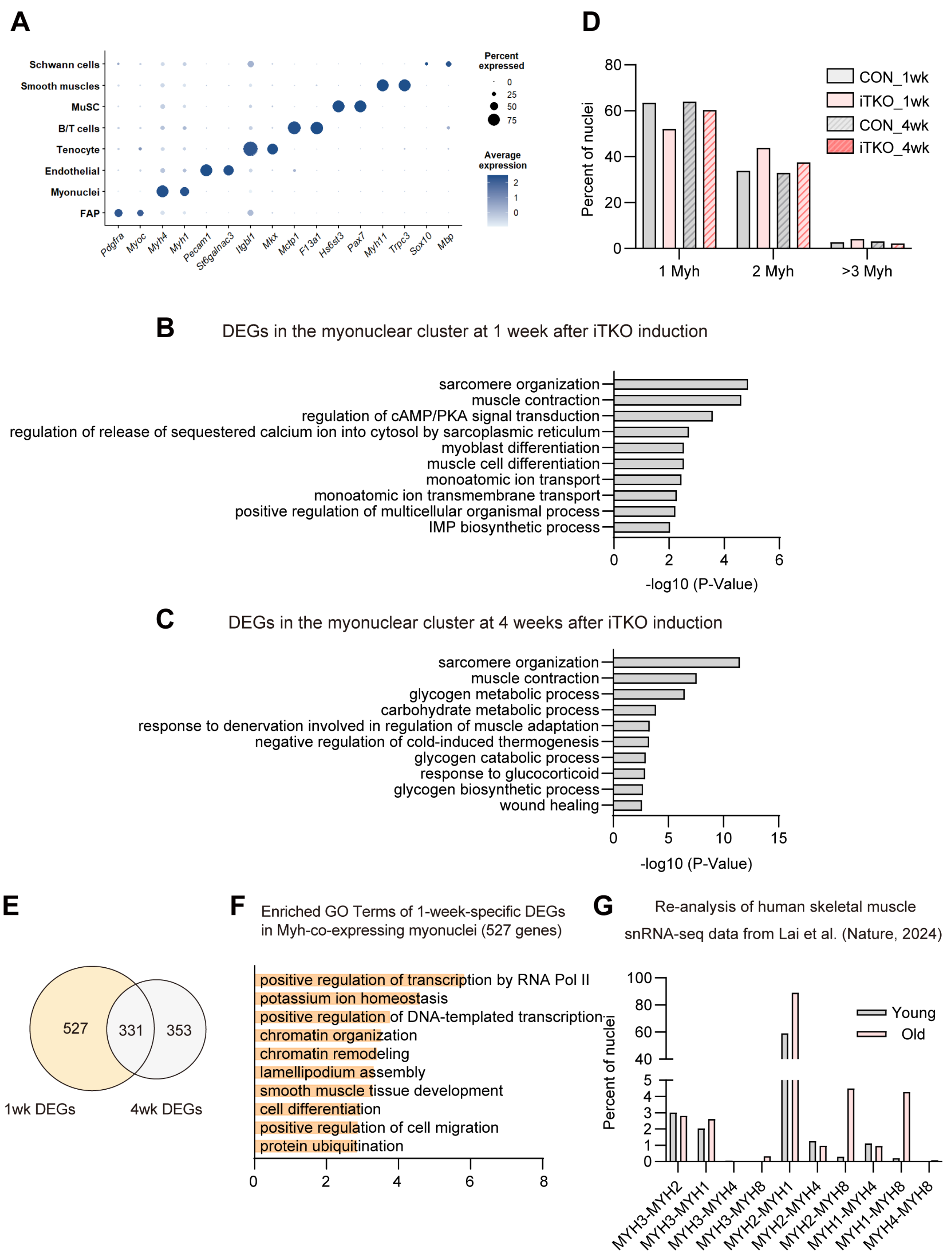


**Figure S7. Time-dependent transcriptional remodeling and increased *Myh* isoform co-expression in large Maf-deficient myonuclei.**

(A) Dot plot showing the expression of representative marker genes used to annotate the major nuclear populations identified by snRNA-seq. Rows indicate nuclear populations, and columns indicate marker genes. Dot size represents the percentage of nuclei expressing each gene, and color indicates the average scaled expression level. (B) Gene Ontology (GO) enrichment analysis of differentially expressed genes in iTKO myonuclei at 1 week after knockout induction. Selected enriched biological process terms are shown. (C) GO enrichment analysis of differentially expressed genes in iTKO myonuclei at 4 weeks after knockout induction. Selected enriched biological process terms are shown. (D) Distribution of myonuclei according to the number of fast *Myh* genes detected per nucleus in control and iTKO muscles at 1 and 4 weeks after knockout induction. Fast *Myh* genes included *Myh2*, *Myh1*, and *Myh4*. Myonuclei were classified as expressing one, two, or three or more fast *Myh* genes. (E) Venn diagram of differentially expressed genes identified in fast *Myh*-co-expressing myonuclei at 1 and 4 weeks after tmx induction. (F) GO analysis of genes selectively altered in fast *Myh*-co-expressing myonuclei at 1 week after knockout induction. (G) Re-analysis of published human skeletal muscle snRNA-seq data showing the proportions of fast myonuclei co-expressing the indicated *MYH* isoform combinations in young and older individuals. Data were obtained from Lai et al. Myonuclei co-expressing *MYH2* and *MYH1*, *MYH1* and *MYH8*, or *MYH2* and *MYH8* are shown.

Supplementary Figure 8 (Sadaki et al.)

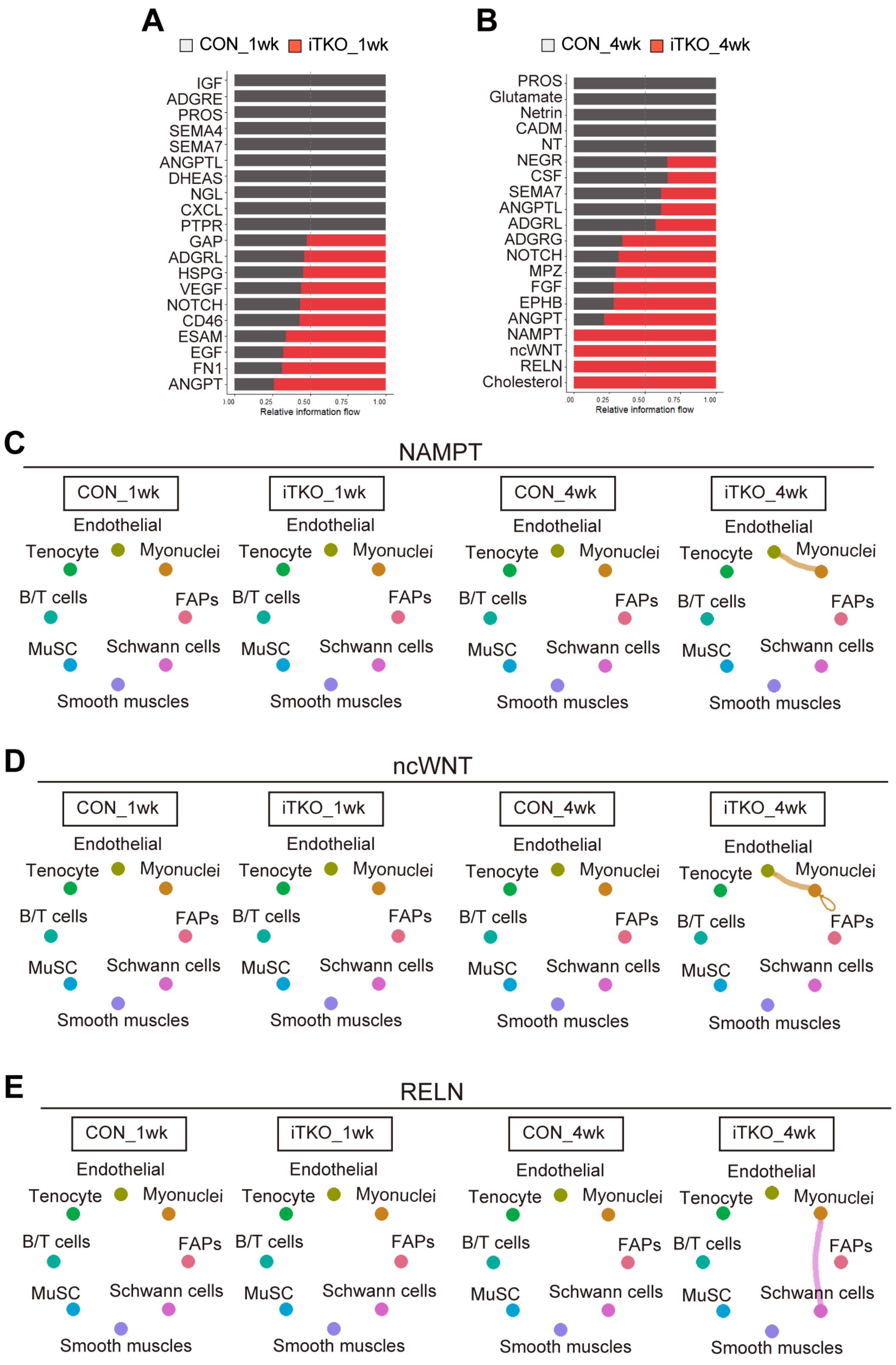

**Figure S8.** Predicted intercellular communication among distinct nuclear populations following large Maf depletion.

(A and B) Relative information flow of the top 20 signaling pathways inferred by CellChat analysis of single-nucleus RNA-sequencing data at 1 week (A) and 4 weeks (B) after knockout induction. For each signaling pathway, stacked bars indicate the relative proportions of the summed interaction probabilities in control and iTKO muscles. (C-E) Predicted intercellular communication among distinct cell populations through NAMPT (C), non-canonical WNT (ncWNT) (D), and RELN (E) signaling at 4 weeks after knockout induction. NAMPT and ncWNT signaling included predicted interactions between myonuclei and endothelial cells, whereas RELN signaling included predicted interactions between myonuclei and Schwann cells. Line thickness represents the predicted interaction strength between cell populations.

**Supplementary Table 1. List of primer sequences.**

| **Gene (genotyping)** | **Orientation** | **Sequence (5' to 3')** |
|---|---|---|
| Mafa | Fw | TATGAGGCCTTCCGGGGTCAGAGCTTC |
| | Rv (WT) | TCTGTTTCAGTCGGATGACCTCCTCCTTGC |
| | Rv (KO) | CGATTGTCTGTTGTGCCCAGTC |
| Mafb flox | Fw | AGGTATGACTGTGTGGCT |
| | Rv | CAAGCCAGAATGCAAAAGCG |
| Maf flox | Fw | AGACCACCTCAAGCAGGAGA |
| | Rv | GTTACGGGGGAATTCAGGTT |
| Cre | Fw | GGACATGTTCAGGGATCGCCAGGCGT |
| | Rv | GCATAACCAGTGAAACAGCATTGCTG |
| oIMR0042 | (Internal control) | CTAGGCCACAGAATTGAAAGATCT |
| oIMR0043 | (Internal control) | GTAGGTGGAAATTCTAGCATCATCC |
| **Gene (RT-qPCR)** | **Orientation** | **Sequence (5' to 3')** |
| *Mafa* | Fw | AGGCCACCACGTGCGCTTGG |
| | Rv | GCTGCTGCACCCGCTTGAAG |
| *Mafb* | Fw | CAACGGTAGTGTGGAGGACC |
| | Rv | CTTCTGCTTCAGGCGGATCA |
| *Maf* | Fw | GCAATGAACAATTCCGACCT |
| | Rv | CCGGTTCCTTTTTCACTTCA |
| *Tbp* | Fw | CAGCCTCAGTACAGCAATCAAC |
| | Rv | TAGGGGTCATAGGAGTCATTGG |
| *Myh4* | Fw | CACCTGGACGATGCTCTCAGA |
| | Rv | GCTCTTGCTCGGCCACTCT |
| *Myh2* | Fw | TGGAGGGTGAGGTAGAGAGTG |
| | Rv | TTGGATAGATTTGTGTTGGATTG |
| *Myh1* | Fw | CGGAGTCAGGTGAATACTCACG |
| | Rv | GAGCATGAGCTAAGGCACTCT |
| *Myh3* | Fw | CAACCCTTATGACTACCCGTTCA |
| | Rv | GATGGCACTATCCGTAGCTAACA |
| *Myh8* | Fw | CAGGAGCAGGAATGATGCTCTGAG |
| | Rv | AGTTCCTCAAACTTTCAGCAGCCAA |
